# A multi-modal fitting approach to construct single-neuron models with patch clamp and high-density microelectrode arrays

**DOI:** 10.1101/2022.08.03.502468

**Authors:** Alessio Paolo Buccino, Tanguy Damart, Julian Bartram, Darshan Mandge, Xiaohan Xue, Mickael Zbili, Tobias Gänswein, Aurélien Jaquier, Vishalini Emmenegger, Henry Markram, Andreas Hierlemann, Werner Van Geit

**Affiliations:** Bio Engineering Laboratory, Department of Biosystems Science and Engineering, ETH Zurich, Basel, Switzerland; Blue Brain Project, École polytechnique fédérale de Lausanne (EPFL), Campus Biotech, 1202 Geneva, Switzerland

**Author notes:** these authors share senior authorship (Andreas Hierlemann, Werner Van Geit).

## Abstract

In computational neuroscience, multicompartment models are among the most biophysically realistic representations of single neurons. Constructing such models usually involves the use of the patch-clamp technique to record somatic voltage signals under different experimental conditions. The experimental data are then used to fit the many parameters of the model. While patching of the soma is currently the gold-standard approach to build multicompartment models, several studies have also evidenced a richness of dynamics in dendritic and axonal sections. Recording from the soma alone makes it hard to observe and correctly parameterize the activity of non-somatic compartments.

In order to provide a richer set of data as input to multicompartment models, we here investigate the combination of somatic patch-clamp recordings with recordings of high-density micro-electrode arrays (HD-MEAs). HD-MEAs enable the observation of extracellular potentials and neural activity of neuronal compartments at sub-cellular resolution.

In this work, we introduce a novel framework to combine patch-clamp and HD-MEA data to construct multicompartment models. We first validate our method on a ground-truth model with known parameters and show that the use of features extracted from extracellular signals, in addition to intracellular ones, yields models enabling better fits than using intracellular features alone. We also demonstrate our procedure using experimental data by constructing cell models from *in vitro* cell cultures.

The proposed multi-modal fitting procedure has the potential to augment the modeling efforts of the computational neuroscience community and to provide the field with neuronal models that are more realistic and can be better validated.

**Author Summary:** Multicompartment models are one of the most biophysically detailed representations of single neurons. The vast majority of these models are built using experimental data from somatic recordings. However, neurons are much more than just their soma and one needs recordings from distal neurites to build an accurate model. In this article, we combine the patch-clamp technique with extracellular high-density microelectrode arrays (HD-MEAs) to compensate this shortcoming. In fact, HD-MEAs readouts allow one to record the neuronal signal in the entire axonal arbor. We show that the proposed multi-modal strategy is superior to the use of patch clamp alone using an existing model as *ground-truth*. Finally, we show an application of this strategy on experimental data from cultured neurons.

## Introduction

In computational neuroscience, multicompartment models provide arguably the one of the most bio-physically detailed representations of single neurons. They are built by combining a precise morphological reconstruction of neurons, obtained through imaging techniques, with electrophysiological characteristics of ion-channel dynamics and their distribution over the neuron morphology.

The use of multicompartment models has enabled researchers to explore several characteristics of neuronal dynamics, including active dendritic properties [39, 5, 29] and the role of the axonal initial segment in initiating action potentials [37, 31, 24]. The models also were used to generate experimentally testable hypotheses to drive research forward. In recent years, we have witnessed massive international efforts in constructing and sharing biophysically detailed multicompartment models. The Neocortical Microcircuit Portal [58, 52] of the Blue Brain Project contains thousands of publicly available cell models of the rat somatosensory cortex. A similar effort is being conducted by the Allen Institute of Brain Science, whose cell-type database includes a growing number of cell models of mice and even humans [35].

Historically, multicompartment models were constructed by taking ionic mechanisms from preexisting knowledge bases and by manually and iteratively tuning the many parameters of the models, until the model matched the *expected* or observed behavior [6]. In more recent years, computer-based optimization has enabled computational neuroscientists to explore the large parameter space more thoroughly and faster, for example by using evolutionary strategies to search for sets of parameter values that provide good fits to experimental data [21].

For the vast majority of available multicompartment models, experimental features used to fit the model are extracted solely from somatic patch-clamp recordings. While the soma clearly is a very important “compartment” of a nerve cell, neurons are much more than just their somata. For example, complex dynamics can arise from active dendritic properties (calcium spikes, non-linear integration etc.) [49, 47, 69, 39] and from the axon initial segment, which is known to be the location where spikes initiate, and which has a determining role for neuron excitability [43, 20]. However, performing simultaneous patch-clamp recordings of multiple compartments of the same cell is extremely challenging and tedious [50, 39, 5].

A strategy to capture neuronal firing dynamics over a larger spatial range includes the use of extracellular recordings. Extracellular signals are generated by transmembrane currents of all neuronal compartments [55, 51, 36] and provide an indirect readout of the intracellular signaling. Gold et al. [32] first suggested that extracellular electrical recordings could provide suitable data for devising extracellular models, but their suggestion has not been followed up by the modeling community. With the advent of high-density micro-electrode arrays (HD-MEAs), however, extracellular signals have been demonstrated to enable the recording, especially *in vitro* (and to a certain extent also *in vivo* [46, 45]), of signal of individual neurons at sub-cellular resolution [53]. The richness of information obtained from extracellular signals enables one to observe electrical potential and signal distribution over the entire neuron. Therefore, extracellular recordings allow for parameterizing neuronal compartments, which cannot be simultaneously and directly probed by patch-clamp experiments.

Here, we present a proof-of-concept study combining patch-clamp and HD-MEA data to construct single-neuron multicompartment models. We first give a general overview of the workflow and describe the steps to include extracellular signals as an extra source of data into the models. Next, we validate our approach on a ground-truth model with known parameters, to assess the benefit of including extracellular data on the final outcome. Finally, we present examples of cell models, constructed from rat cortical cell cultures, and discuss the differences and improvements of our multi-modal strategy over the use of patch clamp alone.

### Workflow to construct multicompartment models from patch-clamp data

Before diving into our multi-modal fitting approach, we introduce here the general workflow for constructing multicompartment models, in order to lay out the different steps and terminology that are used by computational neuroscientists to build biophysically detailed models of single neurons. The main objective of the construction of a multicompartment model is to build an *in silico* replica of a real neuron that can be used to describe the electrical activity of a neuron. The pipeline to construct multicompartment models consists of several steps (Figure 1) that are outlined in the following.

**Figure 1:**
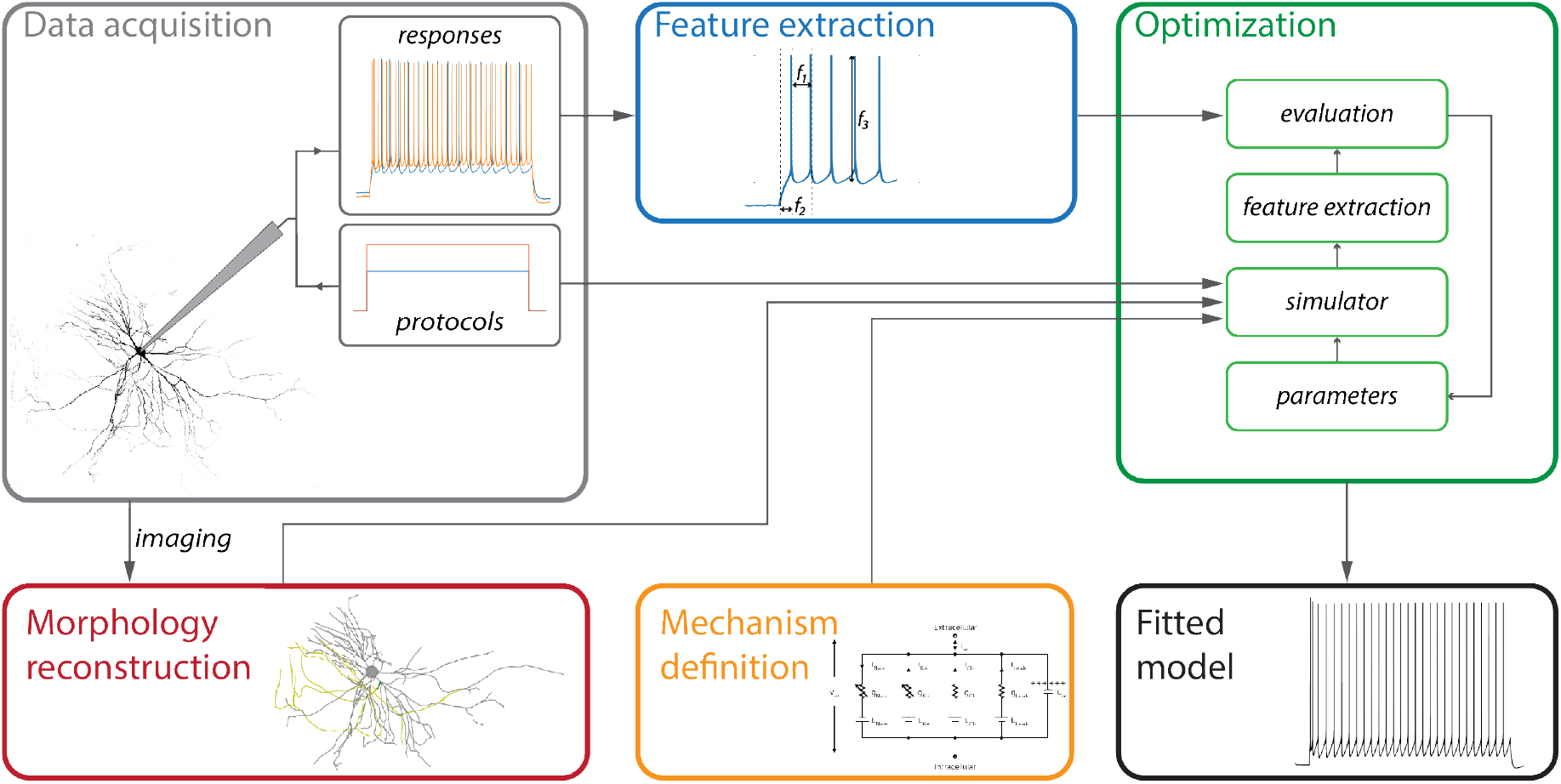
General workflow to construct multicompartment models. In the Data acquisition phase, a patched neuron is stimulated by applying different *protocols*, and the respective *responses* are recorded. Then, the patched neuron is imaged, and the neuron morphology is reconstructed. Mechanisms need to be defined that determine or describe the electrophysiological characteristics of the neuron. The corresponding parameters are then used for the fitting. Features of the measured neuronal characteristics and responses are extracted to reduce the dimensionality of the recorded data. In the Optimization phase, the neuron morphology and the defined mechanisms are used to emulate the applied protocols, and the simulated electrophysiological characteristics are evaluated against the experimentally obtained ones. This procedure is repeated while iteratively varying parameters until convergence is reached. Finally, the morphology, mechanisms, and optimized parameters yield the Fitted model.

#### Data acquisition

the first and essential step is the data collection. A target neuron is patched with a micro-pipette in current-clamp mode to record its **responses** to different stimulation **protocols**. The different protocols are designed to unveil different characteristics of neural dynamics, such as firing patterns, sub-threshold behavior, or spike adaptation. Each protocol is usually applied in several runs. The patch pipette is loaded with a dye (either biocytin or a fluorescent dye) that fills the cytosol of the target neuron during the recording session and that is used for reconstructing the neuron morphology.

#### Morphology reconstruction

after the electrophysiology data acquisition, the target neuron is imaged to reconstruct its 3D morphology. The imaging consists of a z-stack of high-resolution images, which allows for a precise reconstruction of the neurites and their diameters using specialized software tools (e.g. Neurolucida©, Simple Neurite Tracer – SNT [8], Vaa3d [57]).

#### Definition of model mechanisms

the neuron morphology only provides geometrical information for the model. In order to reproduce neuronal behavior, the model needs to be *populated* with **mechanisms**, which govern the equations that give rise to the neuronal dynamics. Mechanisms can be passive (leaky) or active (voltage-gated ion-channels), the latter are usually modeled with a Hodgkin-Huxley formalism [39, 52]. Additional mechanisms can further control the dynamics of specific ion concentrations, for example of Ca^2+^, which can affect the dynamics of sub-populations of ion channels. Different mechanisms are defined in different neuronal sections (e.g., soma, dendrites, axons). Computational neuroscientists need to select the mechanisms for the different neuronal compartments, which will greatly influence the performance of the final model. The definition of the mechanisms also determines the **parameters** that need to be identified during the fitting procedure. Parameters include, for example, the maximum conductances of an ion-channel in a certain compartment; properties of passive components, such as membrane or axial resistances; or free variables that are included in the mechanistic equations, such as the decay of calcium concentrations over time.

#### Feature extraction

As the goal of the model is to optimally reproduce experimental data, one needs to define a cost function to quantify the fit of the obtained solution. While a straightforward approach could be to compute the point-to-point distance between the experimentally obtained values and the *in silico* counterparts, such a procedure has proven to be sub-optimal in several previous studies [21] due to the intrinsic variability of neural responses even to the exact same input. A better and more viable solution includes processing the *raw* neuronal responses and extracting **features** that describe the response behavior in a compact and informative way. For example, if a protocol is designed to characterize the firing properties of a neuron, one could extract features, such as mean firing rate, spike counts, as well as bursting patterns and adaptation indices. Additional features can be used to describe the action potential (AP) waveforms (e.g., AP amplitude, AP duration) or sub-threshold behavior. For each feature, one can extract the average value and the standard deviation over different runs, namely *μ_exp_* and *σ_exp_.* After extracting all the relevant features from the raw neuronal responses, the model is ready to be optimized.

#### Optimization

The optimization step is aimed at finding good sets of parameters to fit the extracted features. By combining the protocols, morphology reconstruction, and mechanisms, a **simulator** runs the experiment for a given set of parameters. Using the simulated responses, features - analogous to those that have been defined in the feature extraction step - are computed and compared to those extracted from the experimental data. For each feature *i*, a score *s^1^* is computed:

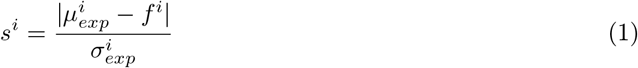

where *f^i^* is the feature value computed from the simulated response, 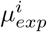 is the mean feature value from the experimental data, and 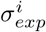 is the experimental standard deviation of the feature. The scores of all features are then summed to compute an overall *fitness* for a certain solution. Several algorithms, mainly based on evolutionary strategies [19], are then used to iteratively explore the parameter space to find solutions with good fitness, i.e., solutions that fit the features extracted from the experimental data well. The optimization procedure yields, upon convergence, the solution (set of parameters) that yields the best fitness. The combination of morphology, mechanisms and the best parameters then constitutes the final multicompartment model.

## Results

### Combining patch-clamp and HD-MEA data for fitting multicompartment models

The general framework described above was developed mainly for use of data from whole-cell patch clamp alone. In order to combine patch-clamp and HD-MEA readouts, we need to augment several steps of the model-fitting pipeline.

First, during the data acquisition, intracellular and extracellular recordings of the target neuron need to be acquired (Figure 2A - left). Therefore, we used cultured neurons from embryonic rats plated on top of an HD-MEA with 26,400 electrodes [53] (see Methods - HD-MEA system for details). The patch-clamp and the HD-MEA systems were synchronized so that we could simultaneously acquire the patch-clamp readout and the extracellular signals from more 500 electrodes in proximity of the patch pipette. Since the HD-MEA signals featured much lower signal-to-noise ratios with respect to the patch-clamp readout, we used the patch-clamp signals to precisely detect action potential times, which we then used to average the extracellular signals (patch-triggered average – PTA). The resulting signal is referred to as an electrical “footprint” (distribution of measured electrical potentials across the array electrodes) or template (Figure 2A - right), and is a clean signal that contains high spatiotemporal resolution information, reflecting the underlying morpho-electric properties of the patched neuron. This footprint or template is used as input data for feature extraction from the extracellular signals. Note that the template is computed using all action potentials acquired during the execution of all protocols, since the more the spikes are acquired, the cleaner the template becomes (extracellular spikes are assumed to be relatively constant across different protocols). In addition, the HD-MEA system we used features a switch-matrix architecture so that not all electrodes but only 1,024 of them are recorded simultaneously [27], which may result in some *holes* in the template. Finally, electrodes recording low-amplitude signals (peak-to-peak amplitude below 5μV) were excluded (Figure 2B).

**Figure 2:**
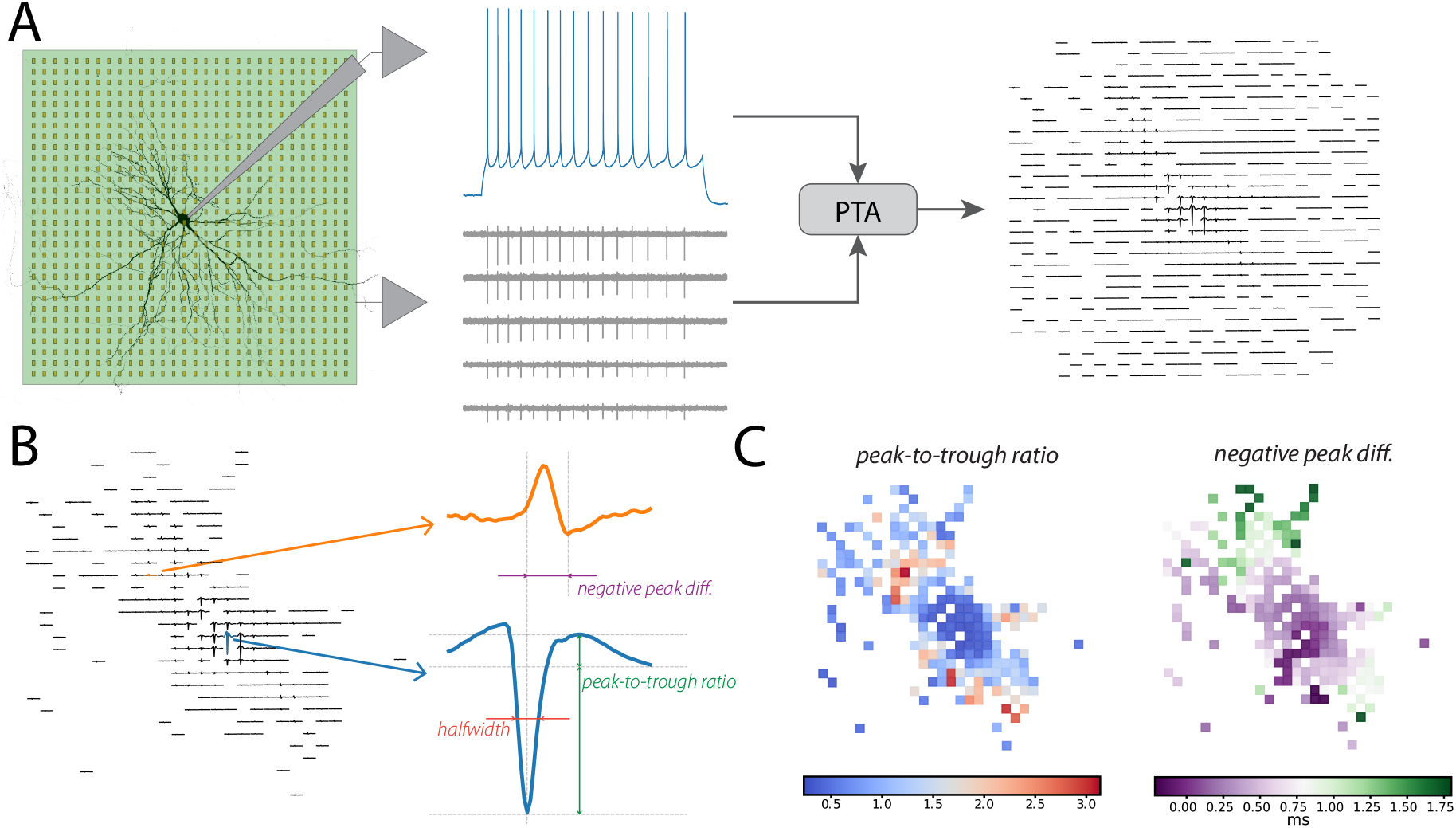
Combining HD-MEA and patch-clamp data. **(A)** Simultaneously recorded patchclamp (blue trace) and HD-MEA data (grey traces) were used to extract the extracellular template using patch-triggered averaging (PTA). **(B)** From the latter (after excluding channels with a peak-to-peak amplitude below 5μV), several extracellular features were computed, either channel by channel (e.g., half-width - red, peak-to-trough ratio - green) or in relation to the channel with the largest signal amplitude (e.g., negative peak time difference - purple). **(C)** Examples of features computed across all the channels: peak-to-trough ratio (left) and negative peak time difference (right).

The second required modification of the overall process was at the feature extraction level. While intracellular features are scalar values, extracted from the somatic patch-clamp trace, e.g., mean frequency or mean AP amplitude, the extracellular template *E* includes *C* electrodes and *T* time points and is, therefore, multi-dimensional in nature (*E* ∈ ℝ^*CxT*^). In order to extract relevant and lower-dimensional features from such templates, we compiled a set of 11 extracellular features *f_ext_* that were defined and used for each recording electrode (or *channel*) (*f_ext_* ∈ ℝ^*C*^). The features either describe channel-specific waveform parameters of the template (termed *absolute* features, such as peak-to-valley duration, peak half-width duration, and peak-to-trough ratio) or values relating the measured value in each channel to that of the channel exhibiting the largest signal amplitude (*relative* features, e.g., negative/positive peak time difference, negative/positive peak amplitude). In total, we defined 11 extracellular features (*N_f_ext__* =11) (see Methods - Extracellular feature extraction for details). In Figure 2B we display the thresholded template (including channels with peak-to-peak amplitude above 5 μV) of a recorded cell, with the largest-signal-amplitude channel depicted in blue and a second channel colored in orange. The right part shows a close-up version of the template for these two channels and some examples of how channel-wise features (peak-to-trough ratio, half-width) were calculated.

The negative peak difference was computed by comparing the negative-peak amplitude of the second channel (orange) with the peak of the largest-amplitude channel (blue). Figure 2C shows the feature maps across the recording electrodes for two exemplary extracellular features, namely peak-to-trough ratio and negative peak time difference.

Still, extracellular features were defined for each channel (*C* can be as high as several hundreds of electrodes), so we explored three different strategies to further reduce the dimensionality of extracellular features that were then included in the optimization:

**single:** with this strategy, a subset of extracellular signals was manually selected (*N_selected_*), and extracellular features were extracted separately for all selected channels. Since the template was obtained by averaging all available extracellular spikes from different runs, the standard deviation of the feature *σ_exp_*, which was used to calculate the feature score (Eq. 1), could not be computed from experimental data. In this case, *σ_exp_* was set to 5 % of the feature value *μ_exp_*. The number of additional extracellular features used for optimization was *N_selected_* · *Nf_ext_*. We used seven single electrodes per cell.
**all:** in order to use the entire information from all extracellular electrodes, we designed a second strategy that used the cosine distance between simulated and experimental features as the score *s_i_*:

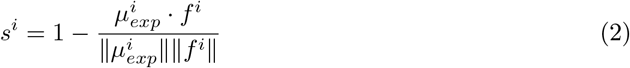

where *i* refers to the *i^th^* feature. This strategy added *N_f_ext__* objectives to the optimization (one scalar for each extracellular feature).
**sections:** the third strategy lies in between the *single* and *all* strategies. In this case, we manually selected *N_sections_* extracellular areas of several electrodes (10-20) that corresponded to different regions of the neuron (e.g., dendritic, perisomatic, axon initial segment areas). For each of these sections, the cosine distance between the simulated and experimental features was used as a score. This strategy provided *N_sections_* · *N_f_ext__* features to the optimization. We used three to four sections per cell.

The different strategies to include extracellular signals allowed us to explore how a modification of the balance between intracellular and extracellular features that were used for the optimization procedure affected the overall fitting performance. Supplementary Figure 1 shows the channel selection for the *single* and *sections* strategies for the different cell models used in the paper.

The final modification that we introduced to the overall pipeline concerned the simulator that was used to generate the simulated responses in the evaluation phase. The `NEURON` software [18] is the gold standard to simulate the intracellular dynamics of single-cell models. With the addition of a Python interface [41], the `NEURON` simulator has been included into the `BluePyOpt` [70] software framework, designed to implement the fitting pipeline, introduced in the previous section, with an easy-to-use and flexible API. In order to simulate extracellular signals in addition to intracellular ones, we extended the `BluePyOpt` framework (see Methods - Extensions to the fitting framework for details). We added a simulator interface based on the `LFPy` [51, 36] Python package. `LFPy` is a wrapper to the `NEURON` software that uses the transmembrane currents from each neuron compartment *I_i_*, obtained via the cable equation, to compute extracellular potentials *ϕ* at the respective electrode positions:

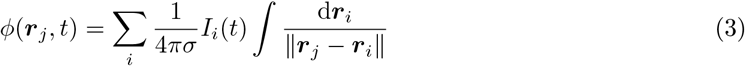

where *σ* is the extracellular conductance (set to 0.3 *S/m* [34]), and *r_i_* and *r_j_* denote the positions of compartment *i* and electrode *j*, respectively. To model the physical size of the recording electrodes, we used the so-called *disk approximation* [36], which averages the extracellular potential over *n* points on the surface of the electrode (we used *n* = 10).

### Validation of the multi-modal fitting strategy on a ground-truth model

After extending the fitting framework to include extracellular signals, we sought to quantitatively assess whether the inclusion of extracellular features improved the overall model fitting performance. To do so, we replicated our planned experiment *in silico*, using an existing multicompartment model with known parameters as ground truth.

The ground-truth model was built using an available morphology of a rat somatosensory cortex layer 5 thick-tufted pyramidal cell (L5PC) [39]. Differently than the original model, where the axon was substituted by a linear *stub* axon of 60 μm, we introduced instead an axon initial segment (AIS) section, expressing AIS-specific conductances [42, 63, 48], namely Na_v_1.2, Na_v_1.6, and K_v_1 (see Methods - Ground-truth model for details). The AIS length was set to 35μm following the reconstructed axon morphology with a subsequent 1-mm long myelinated axonal section with a diameter of 0.2 μm (Figure 3B left). The AIS ion channels in the model moved the initiation of action potentials in the distal part of the AIS (Figure 3B right), in accordance with a large body of experimental and computational evidence [37, 10, 43]. The AIS properties were also strongly reflected in the extracellular signals. The AIS was found to be the dominant contributor to extracellularly measured potentials, due to the strong membrane currents generated to initiate an action potential [66, 10]. The ground-truth model was placed on a 20-row-by-4-column planar simulated MEA with 50 μm electrode-to-electrode center pitch. The extracellular template was computed with Eq. 3 and averaged over several action potentials in response to a step protocol. Figure 3A shows the ground-truth neuron sitting on top of the MEA model. The black lines are the extracellular template at the electrode locations. The red trace depicts the signal with the largest amplitude, which appears on the electrode closest to the AIS. The ground-truth model includes 33 free parameters (10 somatic, 13 AIS, 6 apical, and 4 basal parameters) that need to be optimized. See Methods - Ground-truth model and Table 2-3 for details on the ground-truth model definition and parameters.

**Figure 3:**
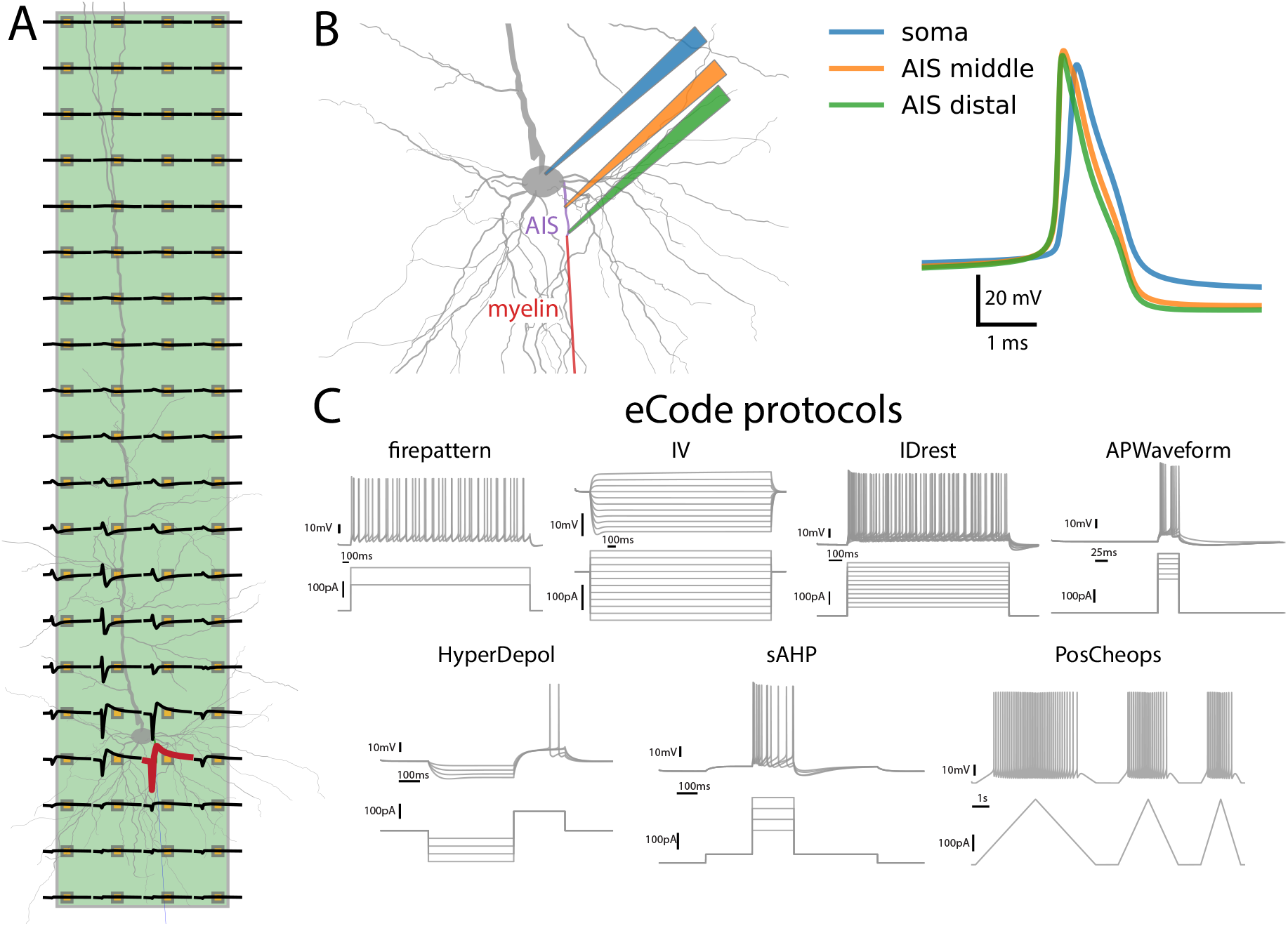
Ground-truth model. **(A)** Visualization of the neuron on top of the 80-channel rectangular MEA used for the simulations. The extracellular electrical signal template is overlaid in black. The red trace depicts the channel with the largest amplitude, which is the channel closest to the AIS section. **(B)** (left) Morphology of the ground-truth neuron, highlighting the additional AIS and myelin sections. (right) Sample action potential signals at the soma (blue), middle AIS (orange), and distal AIS (green). The AP is initiated at the distal end of the AIS and travels back towards the soma. **(C)** eCode protocols used in the *virtual* experiment. For each protocol, the top traces show the somatic membrane potentials and the bottom ones the applied current stimuli.

After checking that our model could reliably reproduce AIS-specific physiological features, which strongly influence the extracellular electrical-potential landscape, we fully replicated the data acquisition phase *in silico*. We utilized a set of protocols named eCode [52, 44] to probe the behavior of a patched neuron against several different stimuli (Figure 3C). The eCode protocols included several long (>1 s) step stimuli with different lengths and amplitudes (*firepattern, IV, IDrest*) to monitor firing behavior, sub-threshold responses, and bursting properties, as well as short-pulse steps (50 ms) to observe details of AP properties (*APWaveform*). Additional more complex stimuli included a hyperpolarizing step, followed by depolarization (*HyperDepol*), a 3-step protocol to monitor After-Hyperpolarization Potentials (AHP) after several spikes (*sAHP*), and, finally, a series of triangular stimuli of different lengths (*PosCheops*). All current amplitudes were computed with respect to the neuron’s rheobase current (estimated using a 270 ms step). For our ground-truth model, the rheobase current was 150 pA. See Methods - eCode protocols for details about amplitudes and timings of the eCode protocols.

From the eCode-protocol ground-truth responses, we then extracted features before running the optimization step. We selected a subset of protocols that were used to fit the model (i.e., training) and we left the other protocols for validation to assess how general the solutions were. For both, the *in silico* and the experimental data we used six protocols for training: IDrest (150%), IDrest (250%), IDrest (300%), APWaveform (290%), IV (−100%), and IV (−20%) (the amplitude is expressed in percentage of the rheobase current).

A total of 76 intracellular features were extracted from these selected protocols and used for optimization assuming that only the patch-clamp data were available (*soma* fitting strategy). In addition to these features, the *all* strategy (using all electrodes and the cosine distance in Eq. 2 as features) adds 11 extracellular features (total=87). The *sections* strategy added 33 (three sections times 11 extracellular features), and the *single* strategy added 77 features (seven electrodes times 11 extra features). The number of intra- and extracellular features for each optimization strategy is summarized in Table 1. See Methods - Feature selection and extraction for details on feature extraction according to different protocols.

**Table 1:** Optimization strategies. Summary of the four different strategies used for fitting the models, specifying the intracellular, extracellular, and total number of features used for optimization (the numbers refer to the ground-truth model).

| strategy name | # intracellular features | # extracellular features | # total features |
| --- | --- | --- | --- |
| *soma* | 76 | - | 76 |
| *single* | 76 | 77 | 153 |
| *all* | 76 | 11 | 87 |
| *sections* | 76 | 33 | 109 |

For each optimization strategy, we ran 10 optimizations, each starting from a different seed using a covariance matrix adaptation (CMA) optimizer (see Methods - Optimization algorithm for details) [19]. Figure 4A shows the fitness of the 10 optimized solutions for each strategy with respect to intracellular (left) and extracellular features (right – features were computed with the *all* strategy for consistency). While the *soma, all*, and *sections* strategies appeared to yield low scores (i.e., better fitness) for intracellular features, the *single* strategy produced solutions with worse intracellular fitness. Conversely, the *single* strategy yielded the best performance with respect to extracellular features, while the worst performance was achieved by using the *soma* strategy. These results reflect the balance between intracellular and extracellular features used for optimization (Table 1). The *single* strategy used almost as many extracellular features as intracellular ones, which may have pushed the optimization to favor extracellular fitting at the expenses of intracellular fitness. In contrast, the use of *all* and *sections* strategies provided a good balance between intracellular and extracellular fitness.

**Figure 4:**
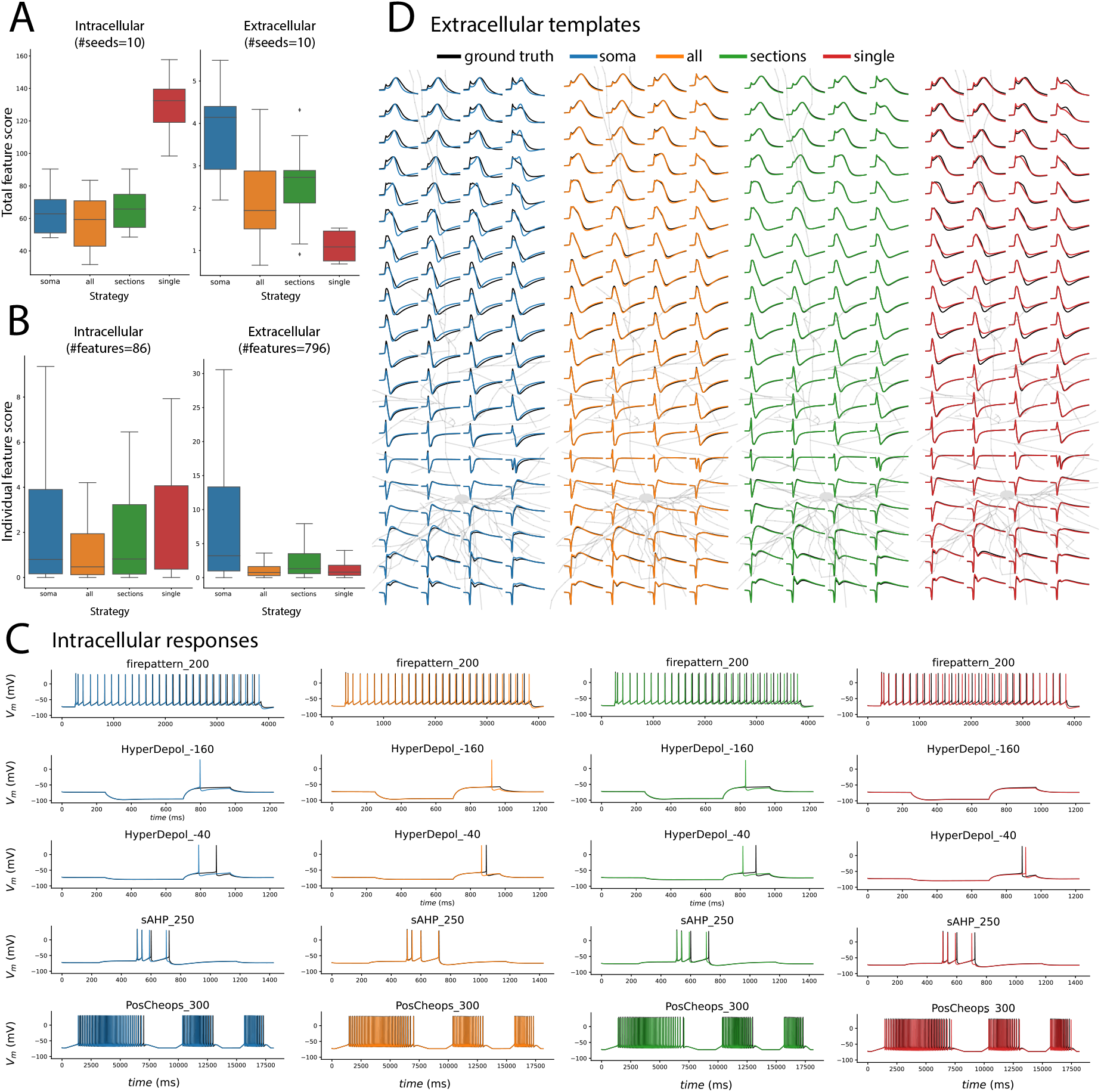
Optimization results on ground-truth model. **(A)** Distributions of intracellular (left) and extracellular (right) scores (i.e., sum of all feature scores) for 10 seeds for each applied strategy (*soma* - blue, *all* - orange, *sections* - green, *single* - red). **(B)** Distributions of intracellular (left, n=86) and extracellular (right, n=796) feature scores for the solution with the best intracellular total score. **(C)** Sample intracellular responses from validation protocols using the seed with the best intracellular score. Black traces are the ground-truth responses. **(D)** Normalized extracellular templates (normalized to the respective template signal-amplitude peak-to-peak value), computed from the firepattern (120%) validation protocol, for the ground-truth model (black) and optimized models (seed with best intracellular score).

Next, we report the results of the solution with the best intracellular fit for each strategy. Figure 4C displays intracellular responses for some validation protocols. One can see that all of these solutions yielded a good fitting of the intracellular traces used for validation (firepattern, HyperDepol, sAHP, and PosCheops). Interestingly, for the HyperDepol stimuli, the *soma, all*, and *sections* strategies failed to reproduce the absence of spikes observed in the depolarization phase (second row – HyperDepol (−160%). Only the *single* strategy correctly did not generate a spike for HyperDepol (−160%), and did generate one for HyperDepol (−40%). The overall distribution of scores for all intracellular features, computed on validation protocols (total=86 features), is shown in Figure 4B-left. No significant differences were detected across strategies (post-hoc Mann-Whitney tests with Holm p-value correction). To quantify extracellular performance, we computed all features using the *single* strategy on all 80 electrodes (total of 796 features after excluding features that resulted in invalid values – e.g., when a clear signal peak was not available). The right panel of Figure 4B shows the distribution of these extracellular features across the different strategies. All strategies making use of extracellular data significantly outperformed the *soma* strategy (post-hoc Mann-Whitney tests with Holm p-value correction: *soma* VS *all* - *p* < 10^-5^, *soma* VS *sections* - *p* < 10^-5^, *soma* VS *single* - *p* < 10^-5^). Figure 4D shows the normalized extracellular templates (each channel value has been normalized to the respective template signal-amplitude peak-to-peak value) for the solution yielding the best extracellular fit. It is evident at first glance that the three extracellular strategies achieve a better fit for almost all electrodes, while the solution from the *soma* strategy, despite providing a very good fit for intracellular features and traces, failed to capture the complex dynamics of extracellular signals in many recording channels.

Extracellular signals indirectly reflect membrane potentials across the entire neuron morphology, as the latter drive transmembrane currents, which are responsible for the generation of extracellular potentials. Having a ground-truth model at our disposal, we could also analyze the intracellular AP distribution, i.e., the AP waveforms across different neurites. We first took the firepattern (120%) response of the ground-truth model and the optimized models, detected action potentials in the somatic trace, and finally used the detected AP peaks to average the action potentials of different neurites (the first and last spikes were excluded as they could contain artefacts originating from stimulus onset and offset). Once the average AP was calculated, we computed the mean relative error of four commonly used features with respect to the ground-truth AP: AP amplitude, AP half-width, decay time, and AHP (afterhyperpolarization). Figure 5 shows the morphology of the ground-truth neuron and the recording positions on the neurites. Each inset displays the AP at the respective location obtained with the ground-truth (black), *soma* (blue), *all* (orange), *sections* (green), and *red single* models (left) alongside with the mean relative errors with respect to the ground-truth AP (right). The selected locations included 4 points along the apical dendrite (apical left, right, middle, proximal), the distal and proximal AIS locations, and two locations on the basal dendrites (basal left, right). For all tested locations, the *all* strategy produces lower errors than the *soma* strategy. For all dendritic locations (both apical and basal), all extracellular strategies (all, *single, sections*) outperform the *soma* strategy. This suggests that combining intracellular and extracellular features could provide a better representation of the AP waveforms across the entire neuronal arbor.

**Figure 5:**
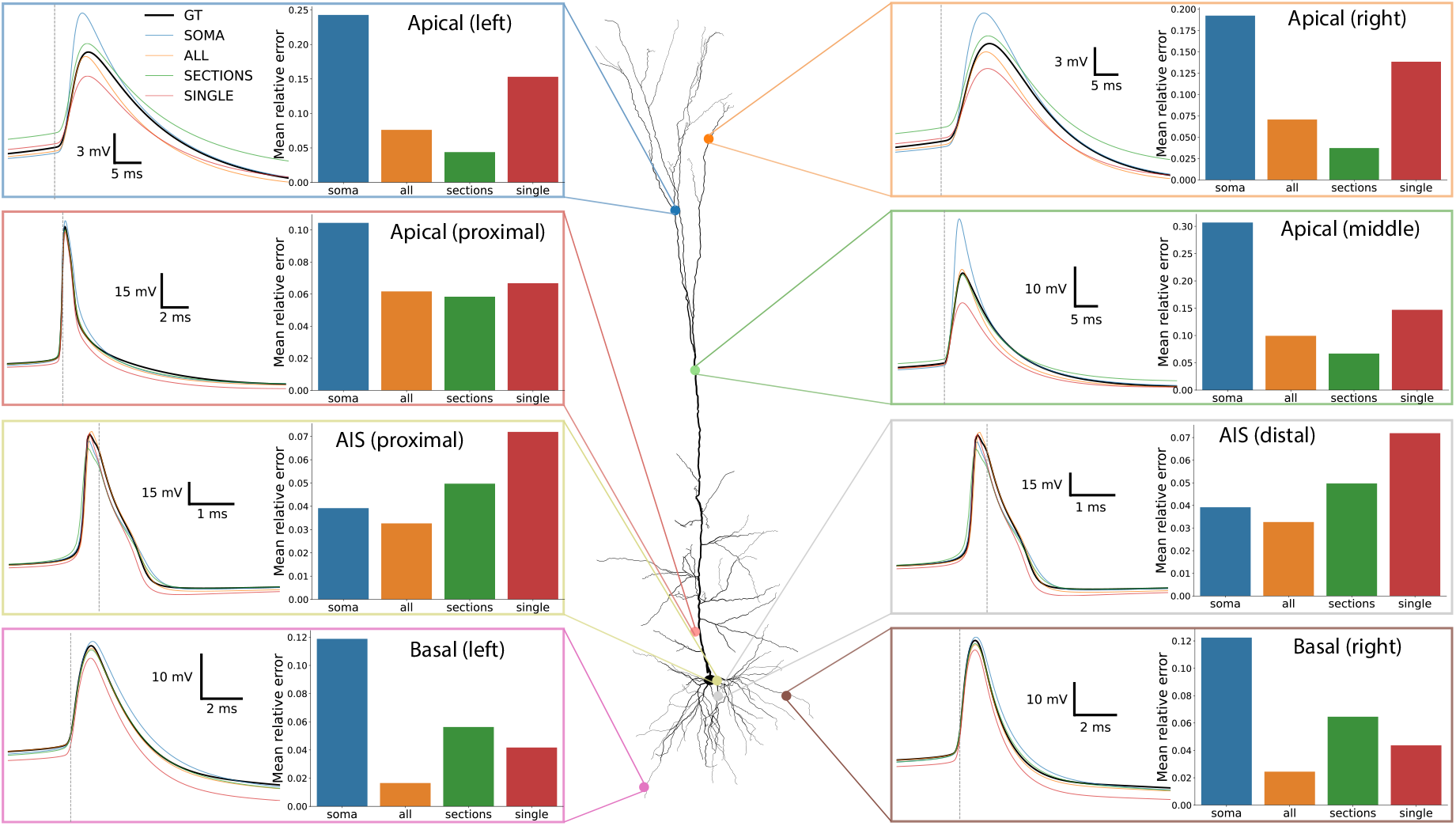
Average AP distribution. Action potentials at different locations along the neuron morphology. Each inset displays the average AP at a specific location on the left (marked on the neuron morphology) and the mean relative error with respect to the ground-truth values of four commonly used features (AP amplitude, AP half-width, decay time, and AHP) for the four different optimization strategies (ground-truth - black, *soma* - blue, *all* - orange, *sections* - green, *single* - red). For the majority of locations, the extracellular strategies yielded the better AP waveforms fits than the *soma* strategy.

In this section, we performed an *in silico* experiment using a ground-truth model with known parameters to investigate whether the inclusion of extracellular features (using different strategies) could be beneficial to construct multicompartment models. We have shown that adding extracellular features, in particular by using the *all* and *sections* strategy, improved the optimization performance, both in terms of fitting of intracellular signals, extracellular templates, and action potential distribution across the entire neuronal morphology. Note that this scenario is ideal because *i)* we imposed that the ground-truth solution lies within the parameter space (i.e., all the mechanisms and respective parameter boundaries are “correct”) and *ii)* the forward model used in the optimization to generate extracellular signals was the same as the one used to generate ground-truth signals.

### Cell models from cultured neurons

After assessing the benefits of combining intra- and extracellular signals with *in silico* experiments, we looked into experimental data. We acquired paired patch-clamp and extracellular signals from two cells in embryonic-rat cortical cultures. After the electrophysiological data acquisition, cells were imaged with a confocal microscope and fixated. We then stained the cultures with AIS-specific anti-bodies to locate the axon initial segment in the morphology. Figure 6 shows the two cells that we used to construct multicompartment models, including a maximum intensity z-projection of the z-stack (top left), the reconstructed morphology overlaid on the extracellular template (top middle – AIS is green, axon is blue, other neurites in grey), sample somatic patch-clamp traces from one of the runs (right), and AIS stainings (bottom - left panel shows Alexa Fluor, the center panel displays the AIS-specific channel yielding the best signal – K_v_7.3 for Cell 1 and Ankyrin-G for Cell 2, and the right panel shows the merged image with an indication of the AIS location). Given the early developmental stage of the cultures (experiments performed between DIVs 18-21), there was no clear and unique apical dendrite. For this reason, when reconstructing the morphology, we did not differentiate between apical and basal dendrite and we applied the same mechanisms to all dendritic sections. Moreover, for both cells the AIS did not originate directly from the soma but from one of the dendrites, which is termed an axon-bearing dendrite (ABD) [67, 30]. Despite these differences, we used the same mechanisms of the ground-truth model detailed in the previous sections and considered the ABD as a “normal” dendrite section (see Methods - Experimental data analysis for details).

**Figure 6:**
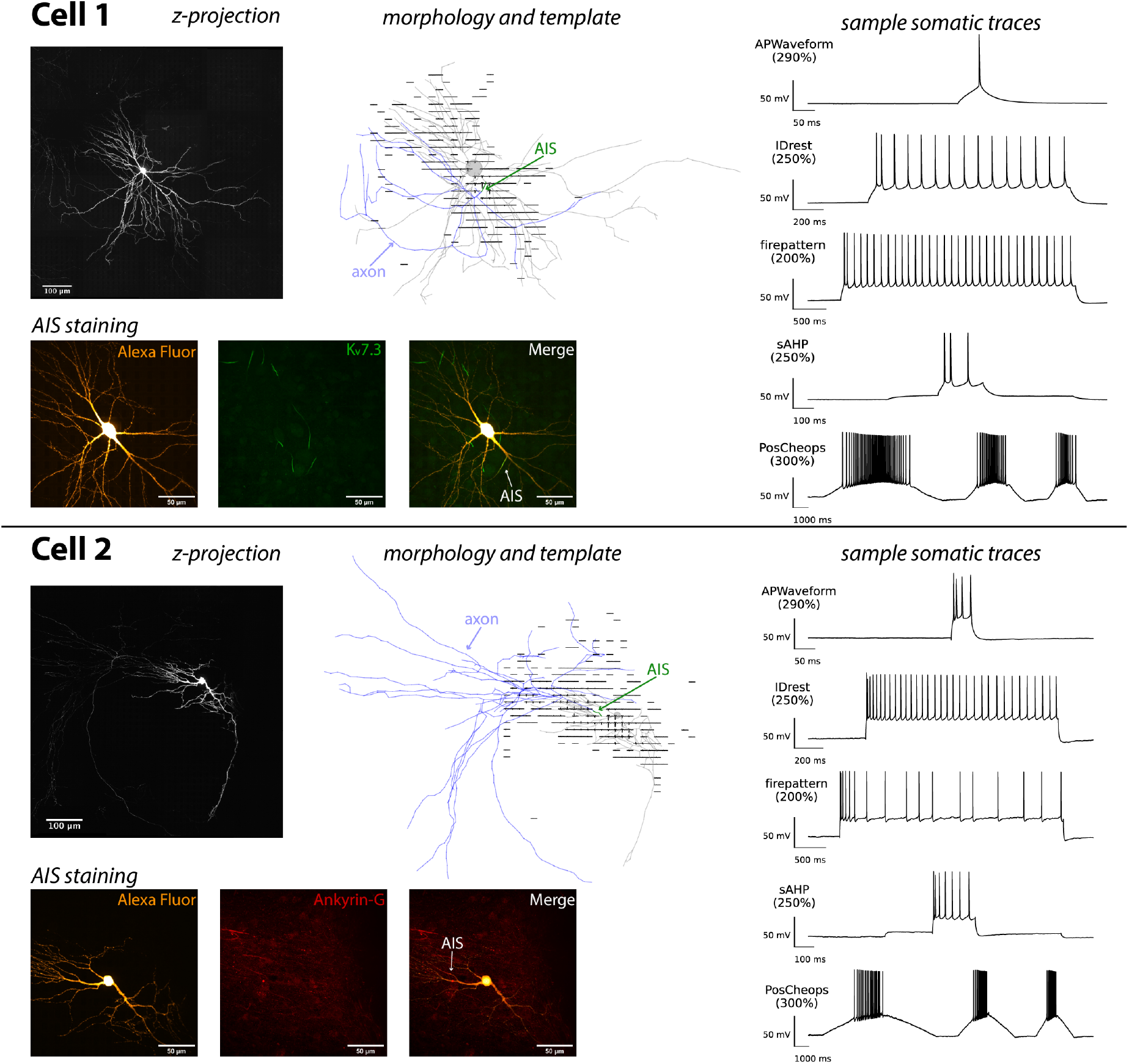
Experimental data. For each recorded neuron (Cell 1 and Cell 2) we display the maximum z-projection of the z-stack (top left), the reconstructed morphology overlaid over the extracellular template (top center – AIS in green, axon in blue), the Alexa Fluor 594 dye and AIS-specific staining (bottom) and sample somatic traces and one experimental run (right).

Figures 7 and 8 show the results of the optimizations performed on Cell 1 and Cell 2 in Figure 6. Panel A displays the intracellular (left) and extracellular (right) scores for the optimized models starting from 10 different seeds for the *soma* (blue) and *all* (orange) strategies (we report results only for the *all* strategy for the extracellular options, as it showed the best performance in the *in silico* tests). For both cell models, the application of the *soma* strategy yielded better intracellular scores, while the use of the *all* strategy resulted in better extracellular fitting. The scores obtained were almost twice the ones we got for the fitting of the ground-truth model in Figure 4 (intracellular score: GT fitting - 62.8 ± 13.6 for *soma* and 59.3 ± 17.0 for *all*; Cell 1 fitting - 123.9 ± 11.2 for *soma* and 149.2 ± 47.2 (excluding one seed that did not converge) for *all*; Cell 2 fitting - 152.8 +- 9.22 for *soma* and 177.6 +- 20.4 for *all*). These discrepancies can be partly explained by the fact that for the *in silico* validation, the optimization procedure was performed with a correct cell model, i.e., by using the same mechanisms and distributions for optimization that have been used the for ground-truth model. On the contrary, for the experimental cells, we did not know the exact ion channels expressed by the neurons and we used the mechanisms present in the ground-truth model. In fact, the experimental cells (putative excitatory neurons recorded in a DIVs 18-24 cell culture) and the ground-truth model (designed to fit a L5PC recorded in a P14-P20 slice) are likely to express different ion channels. Despite these apparent differences, the models fitted on the experimental data could reproduce the main features of the intracellular spiking patterns of the validation protocols (panels B - with the exception of the PosCheops (300%) response for the *soma* strategy in Cell 2 - Figure 8B).

**Figure 7:**
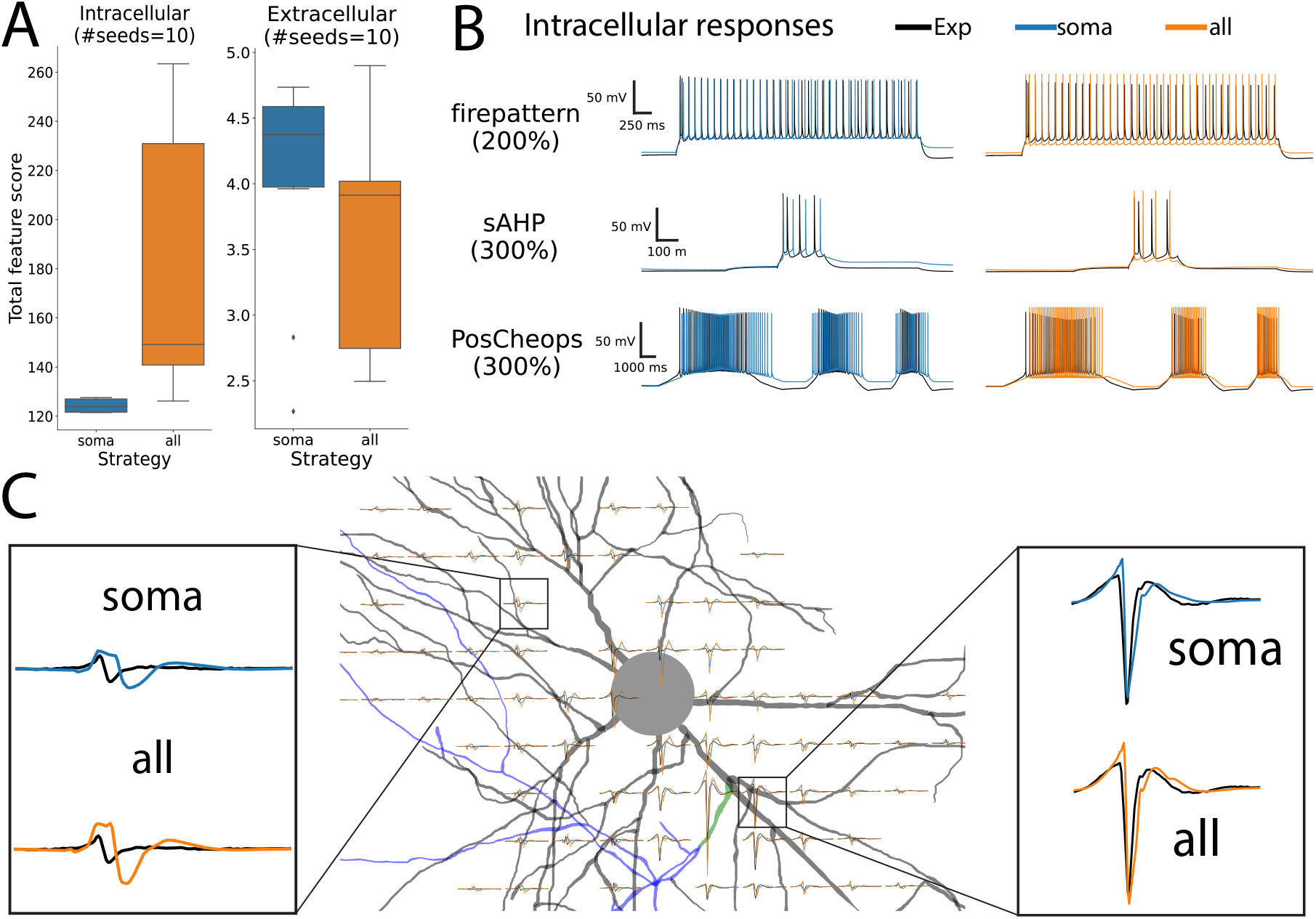
Optimization results for the experimental Cell1 model. **(A)** Distributions of intracellular (left) and extracellular (right) scores (i.e., sum of all feature scores) for 10 seeds for each applied strategy (*soma* - blue, *all* - orange). **(B)** Sample intracellular responses using validation protocols of the seed with the best intracellular score. Black traces show patch-clamp recordings. **(C)** Morphology and normalized extracellular templates, computed using the firepattern (120 %) validation protocol for experimental data (black) and the optimized model (seed with best intracellular score). The insets highlight the experimental (black), *soma* (blue) and *all* (orange) extracellular waveforms on two channels.

**Figure 8:**
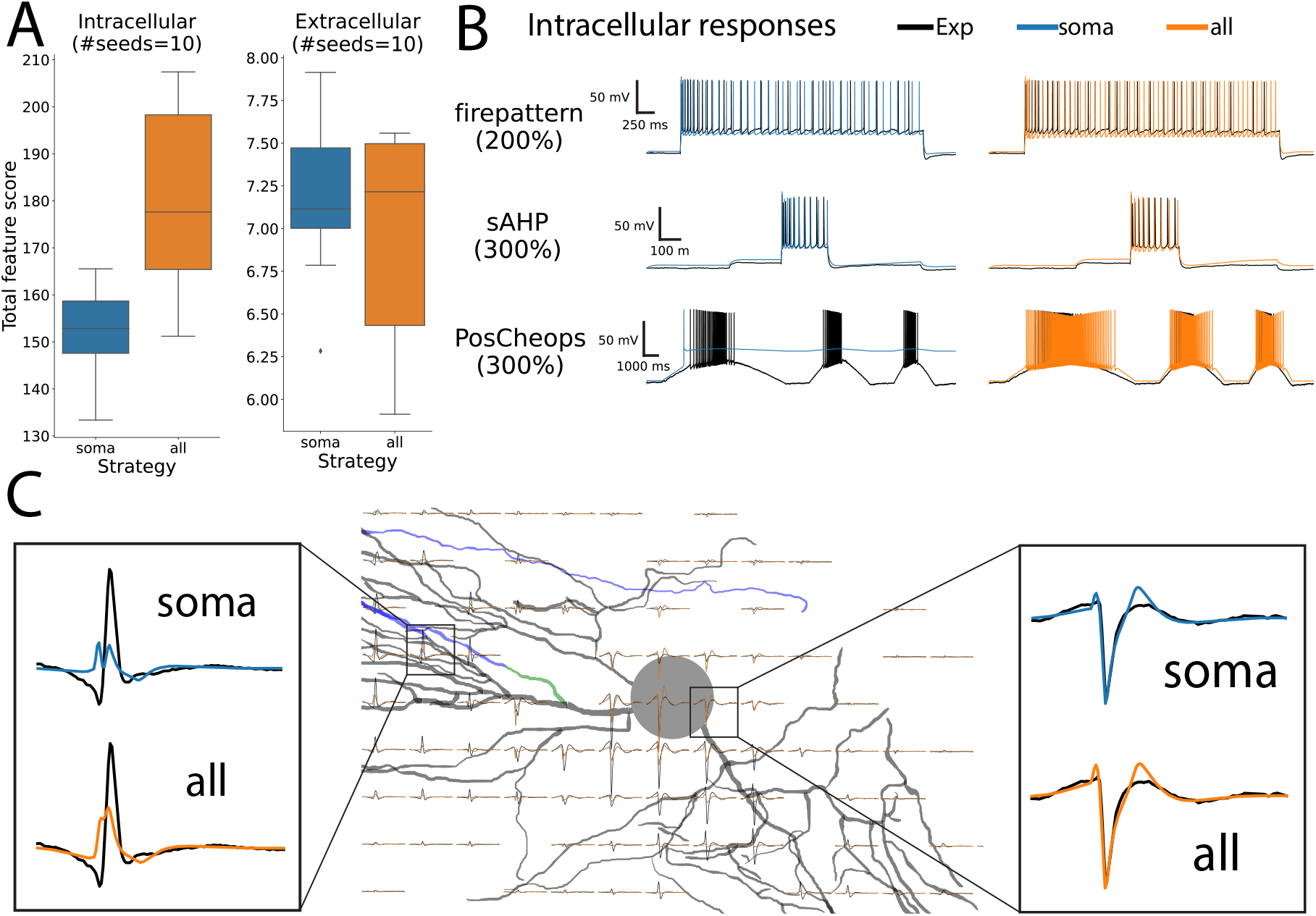
Optimization results on the experimental Cell2 model. **(A)** Distributions of intracellular (left) and extracellular (right) scores (i.e., sum of all feature scores) for 10 seeds for each applied strategy (*soma* - blue, *all* - orange). **(B)** Sample intracellular responses using validation protocols of the seed with the best intracellular score. Black traces show patch-clamp recordings **(C)** Morphology and normalized extracellular templates, computed using the firepattern (120%) validation protocol for experimental data (black) and the optimized model (seed with best intracellular score). The insets highlight the experimental (black), *soma* (blue) and *all* (orange) extracellular waveforms on two channels.

Panels C in Figures 7 and 8 show the experimental extracellular templates (black) and the resulting simulated templates using the *soma* (blue) and *all* (orange) fitting strategies on top of the reconstructed cell morphologies. The insets show a close-up of two representative channels for each cell. The shapes of the fitted extracellular templates of both strategies are qualitatively similar to the experimental templates. The shape characteristics of the waveforms appear to be reproduced by the optimized cell models, but mismatches are still apparent, in particular for Cell 2 (e.g., in the area below the soma in Figure 8). The worse fit of Cell 2 with respect to Cell 1 is also reflected by the overall higher extracellular scores (right boxplots of panels B in Figures 7 and 8).

Considering the confounding factor of probable mismatches between experimentally recorded signal traces and the model traces based on the mechanisms included in the model fits, it is virtually impossible to state whether any of the strategy (*soma* or *all*) is better than the other in this case. Nevertheless, the inclusion of extracellular features in the fitting procedure does not impede the optimization to find acceptable models.

## Discussion

Here, we presented a multi-modal strategy to construct biophysical multicompartment models using a combination of patch clamp and high-density microelectrode arrays (HD-MEAs). After introducing the general concepts of fitting multicompartment models, we presented the extensions that were required to include extracellular signals in the procedure. We then validated the approach using an existing groundtruth model with known parameters and showed that the inclusion of extracellular data improves the overall fitting performance, specifically in terms of fitting extracellular signals and the membrane potentials over the entire neuron morphology. Finally, we applied the validated method to experimental data from two cultured cortical neurons and demonstrated how to build cell models either fitted on the base of patch-clamp data alone or by applying a combined patch-MEA fitting strategy.

While the suggestion to use extracellular data as a source to construct multicompartment models dates back to more than 10 years ago [32], the presented work is the first concise attempt to include high-quality extracellular data into the fitting of single-neuron models. Our *in silico* validation provides convincing evidence that extracellular data can be used to improve the optimization performance and may provide better-validated models. Nevertheless, including extracellular measurements as an additional data source is challenging and may entail various issues that merit discussion. First of all, a setup to perform patch clamp and simultaneous extracellular recordings needs to be available. It requires the use of an upright microscope due to the opaqueness of the MEA substrate. Moreover, the presence of the MEA within the patch-clamp setup may provide additional noise sources for both systems, potentially yielding lower quality data, although several approaches to reduce noise are available. Second, in order to record reliable and large-amplitude extracellular signals, the patched cell needs to be located in close proximity to the extracellular electrodes [17]. The closer the cell, the larger the signal and, therefore, the signal-to-noise ratio. While proximity is not an issue when working with 2D cell cultures where (ideally) cells form a monolayer on top of the MEA, the use of thicker preparations, such as brain slices, may add extra complications: the patch experimenter will need to blindly target *deep* cells that lie close to the surface. However, techniques to use the HD-MEA recordings to estimate the precise patch pipette location help to render the cell targeting easier [4, 56]. Another attractive possibility to use slices is to use organotypic slice cultures [22, 33], where slices are cultured for several weeks on a MEA device and thin down over time. Finally, for imaging the patched neurons, a precise registration between the 3D electrode locations and the neuron morphologies is required in order to correctly assign extracellular signals. Due to the relatively low resolution in the z-dimension of the microscope (in our case 0.4 μm), the electrode registration may include errors that may cause mismatches between experimentally recorded and reconstructed extracellular signals.

The forward-modeling framework used to simulate extracellular signals from transmembrane currents is well-established and widely accepted [23]. However, this framework makes several assumptions that may not be fully satisfied under experimental conditions. The extracellular milieu is assumed to be *infinite*, *isotropic*, *homogeneous*, and *ohmic*. Clearly, the sample is not infinite and it is bounded by the well and the substrate on which the cells are cultured. However, considering that the reference electrodes are at the borders of the sensing area and we targeted cells lying in the center of the HD-MEA [26], the infinity conditions can be considered to be partially met. The isotropy and homogeneity assumption are harder to relax: For *in vitro* cultures, the cells mainly are arranged in a monolayer on top of the substrate. Therefore, the arrangement of cells and medium is not homogeneous and the conductivity of the tissue is not isotropic. Cell density is high and conductivity is low along lateral directions within the cell layer. Cell density becomes lower and conductivity increases upon moving perpendicular to the cell layer into the medium. Moreover, the MEA substrate, on which the cell layer is arranged, is an insulating material, while medium is conductive. However, different analytical solutions can be used to correct for the anisotropy of the tissue [36] and discontinuities, e.g., using the method of images [54], which corrects for the amplitudes of the extracellular potentials. We did not explicitly include these modeling extensions in the forward model, but we did account for the presence of the MEA substrate in computing extracellular features by using relative amplitudes instead of absolute ones. The assumption that the medium shows ohmic behavior seems to be justified in the frequency range of extracellular signals [34]. Besides assumptions concerning the extracellular medium, the cable equation that has been used to calculate transmembrane currents assumes that extracellular potentials outside the cell membrane are constant (i.e., extracellular axial currents are zero), and therefore, self-ephaptic phenomena (the effects of the self-generated extracellular potentials on the neuron itself) are ignored [15]. This assumption is widely accepted due to the very low amplitude of extracellular signals in comparison to intracellular ones. Some of the simplifications and assumptions mentioned above could partially account for the mismatch between simulated and experimentally recorded signals. The use of more advanced and sophisticated modeling frameworks, such as the finite element method [3, 68, 15, 16] could help to overcome some of the simplifications in that MEA substrate and intracellular and extracellular space can be specifically modeled. However, the use of such methods would be computationally very expensive for performing model optimizations with several thousands of simulations until convergence to a stable solution. A probably major contributor to the discrepancies observed between experimental and fitted data (Figures 7 and 8) may originate from the model definition that we used to reconstruct the data of the experimental cells. We used the model structure of a layer 5 pyramidal cell recorded in a P20 slice to fit data recorded of putative pyramidal cells recorded in culture at DIV 18-24. Clearly, the developmental path of neurons in a living animal (from which the slice was obtained) differs from that of cultured neurons in a dish starting from embryonic state. Therefore, the use of an L5PC model to fit neurons of an embryonic cell culture is not ideal. Nevertheless, to the best of our knowledge, there are no models for cell cultured neurons available in literature. As the main goal of this work was not to build models from cultured neurons, but to investigate and develop a methodology and software tools to combine multi-modal data sources in a fitting procedure, the application to experimental data serves only as a proof of concept.

The main intention of introducing the multi-modal method here was to better constrain the many parameters of multicompartment models by using other information-rich data sources. While most of the available models are constructed by using electrophysiological data of a single somatic patch-clamp experiment, a simultaneous patch recording from several neurites, especially dendrites, can shed light on activities and non-linear properties of dendritic regions [50, 39, 5]. Despite being experimentally very challenging, one of the main advantages of such multi-pipette patch techniques is that one can stimulate at both the soma and at the dendrite location to induce calcium spikes and fit the model to reproduce the associated phenomena. While the combined use of a HD-MEA with a somatic patch clamp does not provide intracellular access to dendritic neurites, one can exploit the extracellular stimulation capability of HD-MEAs to selectively stimulate dendrites to induce calcium spikes, the effects of which then can be observed in the soma [39]. Here, we used high-density extracellular micro-electrode arrays as an additional recording modality to probe the spatio-temporal distribution of electrophysiological signals across the entire neuron morphology. However, other multi-modal procedures could potentially constitute attractive alternatives. For example, one could combine the patch-clamp technique with optical electrophysiology readouts to obtain a proxy of the intracellular signals from many different regions of a neuron simultaneously. Genetically encoded voltage indicators (GEVIs) [2] provide indirect membrane potential readout of neuronal compartments that can be used to extract region-specific features for optimization. One drawback of this approach, however, is that neurons would need to be genetically modified to express the fluorescent indicators, which possibly affects electrophysiological characteristics and spontaneous firing of the cell. Considering the low yield of patch-clamp experiments the question arises whether it is possible to build multicompartment neuronal models without patch clamp. The main advantages of the patch-clamp technique include: i) the capability of applying precise stimuli to the neuron through multiple protocols (both supra- and sub-threshold) and ii) filling the neuron with fluorescent markers for subsequent imaging to reconstruct its morphology (which is an essential component of the model). The combination of GEVI imaging and HD-MEA could represent another attractive multi-modal approach that can provide similar features. Electrical stimulation via extracellular electrodes can be used to apply different stimuli to the cell, with the advantage that sub-cellular compartments of the neuron [60] can be targeted; in addition, extracellular stimulation could also be used to generate patch-clamp-like sustained stimuli [1], but probably with lower control and precision than patch-clamp stimulation. The GEVI readout can provide indirect membrane potential measurements across multiple neural compartments, as outlined above, and can also be used to image the target neuron at high resolution for morphology reconstruction owing to the brightness and stability of recently developed indicators [2].

In summary, we developed and investigated a novel approach to combine multiple data acquisition modalities, specifically patch clamp and HD-MEAs, to construct more accurate multicompartment models that can be better validated. As novel neurotechnologies are developed, it can be foreseen that new multi-modal strategies will become available to acquire even richer data sets for building and constraining single-cell multicompartment models.

## Methods

### Extensions to the fitting framework

As part of this project, we extended the open-source BluePyOpt Python framework [70], designed to fit neuroscience models, to enable us to use extracellular signals in the fitting procedure.

### Simulator backend

BluePyOpt originally implemented a `NEURON`-based cell model and simulator (`CellModel` and `NrnSimulator` classes). We implemented two additional classes, namely the `LFPyCellModel` and `LFPySimulator`, that use instead `LFPy` [51, 36] as a backend to construct and simulate neuron models. The `LFPySimulator` can be instantiated with a recording probe object from the `MEAutility` package [13] and computes extracellular signals generated by the neuron’s transmembrane currents (Eq. 3). In addition, a stimulus (`LFPySquarePulse`), response (`TimeLFPResponse`), and recording (`LFPRecording`) classes were added to the framework to support extracellular simulations over the entire pipeline.

### Extracellular feature extraction

We additionally extended the feature-extraction capabilities of `BluePyOpt` to include extracellular features. These features are based on the extracellular template, i.e., the average extracellular action potential. We introduced a new class called `extraFELFeature` that preprocesses the responses to obtain the extracellular template and computes extracellular features from a specified protocol (in our case we used IDrest (300%) – see the eCode protocols section below). Note that in the ideal and noise-free simulation, it is sufficient to use one protocol to compute the extracellular template, while for experimental data we combine several protocols to obtain a cleaner template. The extracellular signals at the respective electrode locations are first interpolated to match the sampling frequency of experimentally recorded extracellular traces (in our case 20 kHz). Optionally, the interpolated traces can also be filtered with a zero-phase filter to mimic the filter applied to experimental traces (in our case we used a Butterworth band-pass filter with cutoff frequencies at 300 and 6000 Hz). The somatic response was used to reliably identify peak times using the `eFEL` “peak_time” feature (the first and last spike can be discarded, because they may contain artefacts of stimulus on- and offsets). Peak times were used to cut-out snippets of the extracellular traces, which were averaged and optionally upsampled to provide more precision in feature values. We used 2 ms before and 5 ms after the peak time as cut-outs and upsampled the template 10 times.

When the template was extracted, extracellular features could be calculated. We compiled a list of 11 features that have been included in the `BluePyOpt` package [7]. Extracellular features are either *absolute*, i.e., computed for each channel separately, or *relative*, i.e., computed with respect to the channel with the largest extracellular signal amplitude:

**peak-to-valley duration** (*absolute*): time in seconds between the negative and positive peaks. If the negative peaks precedes the positive one, the value of the feature is positive. Conversely, when the positive peak precedes the negative one, the value is negative.
**halfwidth** (*absolute*): width of waveform at half of its amplitude in seconds. If the positive peak precedes the negative one, the value is negative (this procedure helps to maximize the shape information carried by the feature value).
**peak-to-trough ratio** (*absolute*): the ratio between positive and negative peaks.
**recovery slope** (*absolute*): after depolarization, the neuron repolarizes until the signal peaks. The recovery slope is the slope of the action potential after the peak, returning to the baseline in dV/dT. The slope is computed within a user-defined window after the peak (default=0.7 ms).
**repolarization slope** (*absolute*): after reaching its maximum depolarization, the neuronal potential will recover. The repolarization slope is defined as the dV/dT of the action potential between the negative peak and the baseline.
**negative peak relative amplitude** (*relative*): the relative amplitude of the negative peak with respect to the negative signal peak of the channel with the largest amplitude. For the largest-amplitude channel, this feature has a value of 1.
**positive peak relative amplitude** (*relative*): the relative amplitude of the negative signal peak with respect to the negative signal peak of the channel with the largest amplitude. For the largest-amplitude channel, this feature has a value of 1.
**negative peak time difference** (*relative*): time difference between the negative signal peak with respect to the negative signal peak of the channel with the largest amplitude. For the largest-amplitude channel, this feature has a value of 1. Note that values can also be negative, if the respective negative signal peak occurs earlier than the negative signal peak on the largest-amplitude channel.
**positive peak time difference** (*relative*): time difference between the positive peak with respect to the occurrence of the positive peak of the channel with the largest amplitude. For the largest-amplitude channel, this feature has a value of 1. Note that values can also be negative if the respective positive signal peak occurs earlier than the positive signal peak on the largest-amplitude channel.
**negative image** (*relative*): voltage values at the time of the negative signal peak on the largest-amplitude channel. The values are normalized by the negative signal-amplitude value on the largest-amplitude channel.
**positive image** (*relative*): voltage values at the time of the positive signal peak on the largest-amplitude channel. The values are normalized by the positive signal-amplitude value on the largest-amplitude channel.

### eCode protocols

The eCode protocols used for both, the *in silico* validation and experimental data acquisition included a series of stimuli designed to probe the neuron’s behavior under a wide range of conditions [52, 44]. After a neuron was patched, the holding current *I_holding_* was manually adjusted so that the read-out membrane potential was −70 mV (which corresponded to ~-84 mV after Liquid Junction Potential correction – see Patch-clamp solutions section below). A series of step stimuli of 270 ms duration with increasing amplitudes were then applied to roughly estimate the neuron’s rheobase current *I_rheo_*, i.e., the current at which a single action potential was induced. The subsequent stimulus amplitudes are reported in percent values of the estimated *I_rheo_* current.

**IDthresh**: the purpose of this protocol was to finely screen the current levels to correctly re-estimate *I_rheo_* before processing. It consisted of 21 sweeps with a 270 ms square pulse of increasing amplitudes from 50% to 130% with step increments of 4%.
**ñrepattern**: this protocol was aimed at characterizing the firing properties of the neuron. It consisted of two step pulses 3.6 s at 120% and 200%.
**IV**: the purpose of this protocol was used to check on sub-threshold properties of the neuron (e.g. the sag). The protocol consisted of 11 step stimuli of 3 s ranging from −140 to 20% with increments of 20%.
**IDrest**: this protocol was used to characterize the input/output curve of the neuron. 11 step stimuli of 1.35 s were delivered with amplitudes ranging from 50% to 300% and increments of 25%.
**APWaveform**: this protocol was defined to precisely record 1-2 action potentials at high sampling rates (typically 200 kHz). It consisted of short 50 ms pulses with amplitudes from 200% to 350% and step increments of 30%.
**HyperDepol**: this protocol was aimed at looking at cell excitability at a hyperpolarization potential. It consisted of 4 sweeps with hyperpolarizing steps of 450 ms from −40% to −160% (increment step 40%), followed by a 270 ms 100% step.
**sAHP**: this protocol was aimed at characterizing After-Hyperpolarization Potentials (AHPs) after several spikes. It consisted of 4 sweeps starting with a 250 ms depolarizing step at 40%, immediately followed by a larger amplitude phase of 225 ms, ranging from 150% to 300% with 50% step increments, and a third 450 ms phase back at 40% amplitude.
**PosCheops**: this final protocol was used to characterize the neuron’s response to ramp stimuli. It contained three ramps of up to 300% amplitude of different duration: the first ramp featured ramp-up and -down phases of 4 s, the second ramp 2 s and the third ramp 1.33 s.

All protocols (excluding IDthres, which was not used for training nor validation) are shown in Figure 3D. For experimental data, the eCode protocols were run 4 to 6 times.

### Ground-truth model

To build the ground truth model, we used the morphology of a previously reconstructed layer-5 pyramidal neuron (L5PC) [39]. The axon was removed except for the first 35 micrometers (axon initial segment or AIS). A 1 mm-long myelinated cylindrical axon with a 0.2μm diameter was attached to the AIS. The model was composed of 4 sections, which contained specific voltage-gated conductances (apical dendrites, basal dendrites, soma, AIS) and 1 section (myelinated axon) that contained only passive channels. The voltage-gated mechanisms were derived from the L5PC model of Markram et al. [52]. In order to improve the simulation of action potential waveforms in the AIS, we replaced the axonal sodium channels present in [52] by two different sodium channels (Na_v_1.2 and Na_v_1.6) that have been shown to be present in the AIS of L5PC [42]. As shown experimentally [42], Na_v_1.2 density gradually decreased, while Na_v_1.6 density gradually increased along the AIS. Moreover, we also added a gradual increase of K_v_1 channels along the AIS [48, 63]. The parameters of the model were optimized in the different sections using BluePyOpt [70] in order to reproduce 58 electrical features of L5PC. The somatic features (e.g., action potential waveforms, firing rates, etc.) were taken from previously recorded L5PC [52]. The dendritic features (action potential backpropagation and calcium entry) were adapted from [64, 40, 62]. The AIS features (action potential waveform) were taken from [59].

### Feature selection and extraction

Relevant electric features (e-features) were extracted from the somatic patch-clamp recordings using the branch BPE2 of the BluePyEfe Python package [11], which itself relies on the eFEL Python package [12] for e-feature computation.

To perform e-feature extraction, the efeatures were first computed on all voltage recordings independently. Then, the rheobase of each cell was computed as the lowest current amplitude triggering at least one spike in the majority of recordings of the cell. Each recording was then associated an amplitude, expressed as a percentage of the rheobase of the cell to which it belongs. Finally, the mean and standard deviation of the e-features were computed across the recordings of same stimulus and amplitude. To perform this operation, a tolerance of 10% was added to the amplitudes. This operation can be better understood as a running average of width 20% of the e-feature value along the amplitude. The end results was a set of e-features where combos e-feature name + stimulus + amplitude were associated to a mean and standard deviation. Note that for the ground-truth model described above, due to its deterministic nature and therefore lack of variance, the efeature standard deviation was set to 5% of the e-feature mean.

Here follows a list of the e-features used as targets for optimization and validation of the models. These e-features were chosen so as to fully describe the firing pattern and shape of the APs seen in the experimental data. As mentioned above, amplitudes were expressed as percentages of the rheobase of the cell. Please refer to the documentation of eFEL for a description of each e-feature.

The E-features used for optimization for different protocols are (total of 79 features):

- **IDrest** (amplitudes: 150%, 250%, 300%): mean_frequency, burst_number, adaptation_index2, ISI_CV, ISI_log_slope, inv_time_to_first_spike, inv_first_ISI, inv_second_ISI, inv_third_ISI, inv_fourth_ISI, inv_fifth_ISI, AP_amplitude, AHP_depth, AHP_time_from_peak, voltage_base, Spikecount (before step), Spikecount (after step)
- **IV** (amplitudes: −100%,-20%): Spikecount, voltage_base, voltage_deflection, voltage_deflection_begin, steady_state_voltage_stimend, ohmic_input_resistance_vb_ssse, decay_time_constant_after_stim, sag_amplitude (only for −100%), sag_ratio1 (only for −100%), sag_ratio2 (only for −100%)
- **APWaveform** (amplitude: 290%): AP_amplitude, AP1_amp, AP2_amp, AP_duration_half_width, AP_begin_width, AP_begin_voltage, AHP_depth, AHP_time_from_peak

The e-features used for validations for different protocols are (total of 110 features):

- **firepattern** (amplitudes: 120%, 200%): mean_frequency, burst_number, adaptation_index2, ISI_CV, ISI_log_slope, inv_time_to_first_spike, inv_first_ISI, inv_second_ISI, inv_third_ISI, inv_fourth_ISI, inv_fifth_ISI, AP_amplitude, AHP_depth, AHP_time_from_peak
- **HyperDepol** (amplitudes: −160%,-120%,-80%,-40%): Spikecount, burst_number, AP_amplitude, ISI_values for the depolarized phase of the stimulus and sag_amplitude, sag_ratio1, sag_ratio2 for the hyperpolarized phase.
- **sAHP** (amplitudes: 150%, 200%, 250%, 300%): Spikecount, AP_amplitude, ISI_values, AHP_depth, AHP_depth_abs, AHP_time_from_peak, steady_state_voltage_stimend
- **Poscheops** (amplitude: 300%): Spikecount, mean_frequency, burst_number, adaptation_index2, ISI_CV, ISI_log_slope, inv_time_to_first_spike, inv_first_ISI, inv_second_ISI, inv_third_ISI, inv_fourth_ISI, inv_fifth_ISI (computed during the three pyramids independently).

### Optimization algorithm

The optimization was performed using the BluePyOpt python package [70] whose optimization module relies on the DEAP Python package [25]. We extended the package and implemented an hybrid CMA optimization strategy [38, 19] tasked to both:

- minimize the sum of the scores defined as 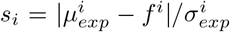, where 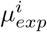 and 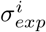 were the experimental mean and standard deviation for e-feature *i*, respectively, and *f^i^* was the feature value computed on the model to evaluate.
- maximize the hyper-volume of the Pareto front formed by the current population of models [9]. At each generation, all models in the population of size λ = 20 were ranked for both criteria, and a mixed rank was obtained following the formula

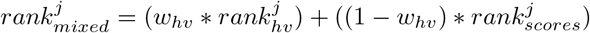

The weight assigned to the hyper-volume ranking, *w_hv_*, was set to 0.4 in the present study. Following this ranking, the *μ* = λ/2 first individuals were selected to update the CMA kernel for the next generation. Note that during the computation of the scores, the scores computed for the extracellular e-features by cosine similarity (all and *sections* strategies) were weighed by a an empirically-obtained factor of 2.5, compared to the intracellular ones, in order to balance the values of intracellular and extracellular scores.

For each model, we ran 10 optimizations of 600 generations, each for a different starting seed of the random number generator.

### Experimental data acquisition

#### HD-MEA system

We used an in-house developed high-density microelectrode array (HD-MEA) system with 26,400 electrodes covering a sensing area of around 2 x 4 mm^2^ [27, 53]. The center-to-center electrode distance was 17.5 μm, and the electrodes were coated with Pt-black to lower impedance and improve the signal-to-noise ratio. The HD-MEA system can be configured to record from up to 1,024 electrodes simultaneously at 20 kHz. Electrode configurations were chosen according to the position of the target neuron.

#### Embryonic rat cortical cultures

All experimental protocols involving animals were approved by the Basel-Stadt veterinary office according to Swiss federal laws on animal welfare, and were carried out in accordance with the approved guidelines. We used rat primary neurons, obtained from dissociated cortices of Wistar rats at embryonic day 18, following the protocol described in Ronchi et al. [60].

Before cell plating, HD-MEA chips were sterilized with 70% ethanol for 30 min. After removing ethanol, the chips were rinsed three times with sterile tissue-culture-grade water. The HD-MEA chips were then coated with a layer of 0.05% polyethylenimine (Sigma-Aldrich, Buchs, Switzerland) in borate buffer to make the surface more hydrophilic. Next, we added a layer of laminin (Sigma-Aldrich, Buchs, Switzerland, 0.02 mg/mL) in Neurobasal medium (Thermo Fisher Scientific, Waltham, Massachusetts, United States) on the HD-MEA and incubated for 30 min at 37°C to facilitate cell adhesion. We used trypsin with 0.25% EDTA (Gibco, Thermo Fisher Scientific, Waltham, Massachusetts, United States), followed by trituration, to dissociate embryonic cortices of E-18 Wistar rats enzymatically and then seeded 15,000 to 20,000 cells in 7 *μ*L on top of the electrode arrays. The plated chips were incubated for 30 min at 37°C before adding 2 mL of plating medium. The plating medium consisted of Neurobasal, supplemented with 10% horse serum (HyClone, Thermo Fisher Scientific, Waltham, Massachusetts, United States), 0.5 mM Glutamax (Invitrogen, Thermo Fisher Scientific, Waltham, Massachusetts, United States), and 2% B-27 (Invitrogen, Thermo Fisher Scientific, Waltham, Massachusetts, United States). After 3 days, 50% of the plating medium were replaced by the BrainPhys™ Neuronal Medium (Stem Cell Technologies) growth medium. The procedure was repeated twice a week. The chips were kept inside an incubator at 37°C and 5% CO_2_. All experiments were conducted between days *in vitro* (DIVs) 18 and 24.

#### Patch-clamp solutions

The intracellular patch-clamp solution consisted of Potassium-gluconate (110 mM), Phosphocreatine (10 mM), KCl (10 mM), Hepes (10 mM), GTP (0.3 mM), ATP-Mg (4 mM) dissolved in nanopure water. Chemicals purchased from Sigma-Aldrich, Buchs, Switzerland. The pH was adjusted to 7.2-7.3 by adding potassium hydroxide at 5 M. On each experimental day, an aliquot was thawed, Alexa Fluor 594 (50 *μ*M) (Sigma-Aldrich, Buchs, Switzerland) was added, and the final solution was filtered using a Millex GV 0.22 *μ*m filter.

The extracellular artificial cerebrospinal fluid (aCSF) solution contained: NaCl (125 mM), KCl (2.5 mM), MgCl_2_·6H_2_O (1 mM), NaH_2_PO_4_ (1.25 mM) and CaCl_2_·2H_2_O (2 mM). On the day of the experiment, 2 L of 1× aCSF was prepared by combining 200 mL of a 10× stock solution with 1 L nanopure water, dissolving 9 g of glucose and 4.2 g of NaHCO3, and topping up to 2 L with nanopure water. This solution was bubbled with carbogen (5% CO2 / 95% O2) and heated to a temperature of around 34°C throughout the experiment.

The combination of intra- and extracellular solutions yielded a Liquid Junction Potential (LJP) of ~ 14 *mV* that we used to correct the values of the recorded membrane potentials.

#### Simultaneous HD-MEA and patch-clamp recording

The experiments were conducted with a custom patch-clamp rig with an upright microscope equipped with the HD-MEA recording unit. The patch-clamp system included an MultiClamp 700B amplifier (Axon Instruments, Victoria, Australia), Axon Digidata 1440A (Axon Instruments, Victoria, Australia), PatchStar Micromanipulator (Scientifica, Uckfield, United Kingdom) and an Olympus BX61WI microscope (Olympus, Tokyo, Japan). The WinWCP software was used to control the patch-clamp system and acquire the intracellular recordings. Glass micropipettes (4-8 MΩ) were pulled using a P-1000 micropipette puller (Sutter Instruments, Novato, California, United States). Whole-cell current-clamp recordings were low-pass filtered at 10 kHz and digitized at 20 kHz. After whole-cell current-clamp mode was established with a target cell, the holding current was manually adjusted to obtain a resting membrane potential of around −70 mV (note that with LJP correction this corresponds to −84 mV). Brief current pulses to induce action potentials were then delivered to the neuron in order to assess that also the extracellular signals reliably showed spikes. We discarded cells with extracellular spikes lower than 50 μV. The patch-clamp and HD-MEA systems were synchronized by sending TTL pulses from the patch-clamp output channel to the field-programmable-gate-array (FPGA) board controlling the HD-MEA device at the beginning of each sweep. Next, we roughly estimated the patched neuron’s rheobase current using increasing step pulses of 270 ms duration. The estimated rheobase current was used to automatically generate the eCode protocol files that were then injected into the neuron. The eCode protocols were repeated during 4-6 runs (each run lasted ~ 200 s).

#### Confocal imaging

After the electrophysiology data acquisition, the chip was mounted under a Nikon NiE upright microscope, equipped with a Yokogawa W1 spinning disk scan head, an ORCA-Flash4.0 V2 Digital CMOS camera (Hamamatsu Photonics, Tokyo, Japan), and a 60x/1.00 NA water-objectives (Nikon, Tokyo, Japan). Prior to imaging, we co-registered the location of the electrodes to the microscope stage position using the stage positions of three electrodes at the vertices of the sensing area. The confocal channel for imaging had an excitation laser of 561 *nm* and an emission filter of 609/54 *nm.* The patched neuron(s) were then imaged using multiple tiles covering their entire morphology. For each tile, a z-stack with 0.4 μm z-step was acquired. The x-y resolution of the images was 112.5 nm. We acquired 6 x 4 tiles and 66 z images for cell 1, and 7 x 4 tiles and 61 z images for cell 2.

#### Fixation and staining

After imaging, the culture was fixated by removing the remaining medium and adding 1 mL of a 4% paraformaldehyde (PFA, Thermo Fisher Scientific, Waltham, Massachusetts, United States) solution for 10 min. Subsequently, the chip was washed three times with PBS for 10 min with gentle agitation. Permeabilization of the membrane and blocking of unspecific binding sites were performed in one step by adding 0.5% Triton X-100 (Sigma-Aldrich, Buchs, Switzerland) to the blocking solution (PBS - Thermo Fisher Scientific, Waltham, Massachusetts, United States) with 10% normal donkey serum (Jackson ImmunoResearch, West Grove, Pennsylvania, United States), 0.02% Na-Az (Sigma-Aldrich, Buchs, Switzerland), 1% bovine serum albumin (Sigma-Aldrich, Buchs, Switzerland) for 30 min. The cultures were then stained for AIS-specific antibodies (Ankyrin-G and K_v_7.3) to identify the AIS tract for the morphology reconstruction. The primary antibodies were diluted in the reaction solution (PBS with 3% normal donkey serum, 0.02% Na-Az, 1% bovine serum albumin, 0.05% Triton X-100) and added to the culture. After 2 h at room temperature on a shaker, the primary antibody solution was aspirated, and the culture was washed thrice with PBS. The secondary antibodies were diluted in fresh reaction solution and added to the culture. After 4 h at room temperature on a shaker, the secondary antibody solution was removed, and the culture was washed thrice with PBS. Until imaging, the culture was kept in PBS at 4°C. The antibodies used in this study included: rabbit-*α*-K_v_7.3 (Alomone Labs, Jerusalem, Israel), guinea pig-*α*-AnkG (Synaptic Systems, Cat. No. 386 004), donkey-*α*-rabbit Alexa Fluor Plus 488 (Thermo Fisher Scientific, Waltham, Massachusetts, United States), donkey-*α*-guinea pig Alexa Fluor 647 (Jackson ImmunoResearch, West Grove, Pennsylvania, United States).

### Experimental data analysis

#### Electrophysiology

The combined patch-clamp and HD-MEA signals were analyzed using `SpikeInterface` [14] and NEO [28] to interface with extracellular and intracellular recordings, respectively.

The patch-clamp and extracellular recordings for each run were processed separately. The TTL pulses, sent by the patch-clamp system to the HD-MEA FPGA, were detected in both data streams and used to synchronize the intra- and extracellular traces (all patch-clamp protocols for each run were concatenated at this time). We removed extracellular channels that displayed saturation of the amplifiers (denoising) and filtered the signals with a 3^rd^-order Butterworth filter with cutoff frequencies at 300 and 6000 Hz. Next, the patch-triggered average was used to compute extracellular templates: intracellular action potentials were detected from the patch signals (the PosCheops protocol was discarded because of the strong spike waveform modulation due to the high-frequency firing rate) and used to extract extracellular waveforms, which were further centered around the extracellular peaks on the channel with the largest signal amplitude to correct for possible mismatches between the intra- and extracellular peak times. To obtain a cleaner template, we discarded waveforms, whose amplitudes were over ±2 standard deviations away from the median amplitude on the channel featuring the largest signal amplitude. The template was then computed as the sample-wise median of the waveforms (we used median because it is less sensitive to noise outliers). Templates were extracted for each run, and the template, which yielded the largest number of channels after denoising, was used for downstream analyses.

#### Imaging and morphology reconstruction

We employed Huygens Professional (version 21.10, Scientific Volume Imaging, The Netherlands, http://svi.nl) to perform the image deconvolution and stitching. Specifically, we first deconvolved images using the CMLE algorithm, with SNR:12 and 40 iterations. Subsequently, the deconvolved images were stitched together with an overlap of 10%, using the circular vignetting correction model.

We used the SNT plugin in Fiji [8] to reconstruct the 3D morphology of the neurons. The soma position was reconstructed as a single point by merging all the branches originating from it to the same root. Neurites were tagged as “soma”, “dend” and “axon”. The AIS tract was identified using the staining images and it was labeled as “apic”, since the SWC format that we used to export the morphology did not formally support the axon initial segment tag. Subsequently we re-labeled the AIS correctly at instantiation in BluePyOpt. The radii of the reconstructed paths were fitted using the “Refine/Fit” tool and exported to SWC format. The raw morphology was preprocessed with a custom Python function (https://github.com/alejoe91/multimodalfitting/blob/master/multimodalfitting/imaging_tools/correct_swc.py) that interpolated missing radii (below or equal to 0.1 μm) and smoothed the radii of each path with a 15-sample moving average. The imaging data were also used to estimate the distance between the center of the soma and the underlying HD-MEA plane.

#### Experimental model definition and feature selection

The models based on experimental data were built using the same set of mechanisms and boundaries used for the ground-truth model (Tables 2 and 3). However, due to the early-development stage of the cultured neurons and the absence of clearly defined basal and apical dendrites, all dendrites were considered the same and expressed all the basal mechanisms (no apical dendrites were defined). The axon-bearing dendrite (ABD) was also treated as a basal dendrite. Furthermore, an axonal section was added to the model with the following mechanisms (and same boundaries reported in Table 2 and 3): gNa16bar_Na16, gkbar_Kd, gCa_LVAstbar_Ca_LVAst, gCa_HVAbar_Ca_HVA2, gCa_LVAstbar_Ca_LVAst, gCa_HVAbar_Ca_HVA2, gSKv3_1bar_SKv3_1, gSK_E2bar_SK_E2, gK_Tstbar_K_Tst, gK_Pstbar_K_Pst, gNap_Et2bar_Nap_Et2, decay_CaDynamics_DC0, gamma_CaDynamics_DC0. The exponential decay value of the gNaTgbar_NaTg basal mechanism (see Table 2) was also added as a free parameter, with boundaries of [-0.1, 0]. In total, the experimental models had 42 free parameters to optimize.

**Table 2:**
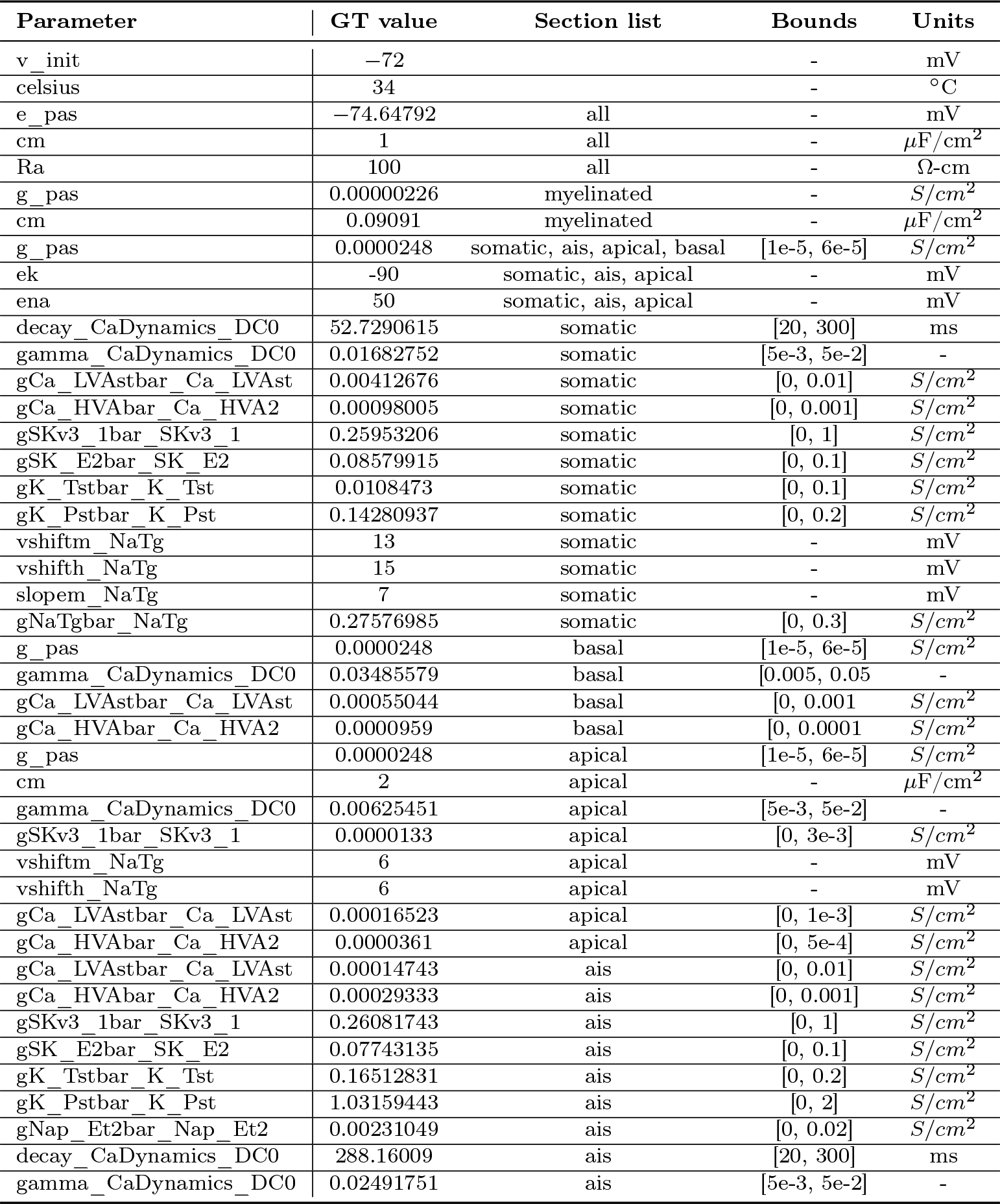
Ground-truth model parameters (non-distribution type) and optimization bounds. Only parameters with optimization bounds are fitted.

**Table 3:** Ground-truth model parameters with distributions and optimization bounds. Only parameters with optimization bounds are fitted.

| Parameter | Distribution | GT value | Section list | Ref. | Bounds | Units |
| --- | --- | --- | --- | --- | --- | --- |
| gNa12bar_Na12 | $\frac{1}{1+\exp\left(\frac{\text{distance}-25}{3}\right)} * \text{Value}$ | 6.15808 | ais | ais[0](0) | [0, 8] | $S/cm^2$ |
| gNa16bar_Na16 | $\left(1 - \frac{1}{1+\exp\left(\frac{\text{distance}-20}{2.5}\right)}\right) * \text{Value}$ | 4.39817 | ais | ais[0](0) | [0, 8] | $S/cm^2$ |
| gkbar_Kd | $\left(1 - \frac{1}{1+\exp\left(\frac{\text{distance}-20}{2.5}\right)}\right) * \text{Value}$ | 0.04356 | ais | ais[0](0) | [0, 2] | $S/cm^2$ |
| gIhbar_Ih | $(-0.8696 + 2.087 * \exp((\text{distance} * 0.0031)) * \text{Value}$ | 0.00001 | somatic,<br>apical, basal | soma(0.5) | - | $S/cm^2$ |
| gNaTgbar_NaTg | $\exp(\text{distance} * (-0.00352)) * \text{Value}$ | 0.06642 | apical | soma(0.5) | [0, 0.1] | $S/cm^2$ |

## Data and code availablilty

All the data and code used in this article are openly available. The contributions to the BluePyOpt – https://github.com/BlueBrain/BluePyOpt – package are summarized in the pull request https://github.com/BlueBrain/BluePyOpt/pull/385 and will be included in the next release of the package. The script, notebooks, and pre-processed data used for optimizations (including cell morphologies, feature and protocol files, extracellular templates and probe files) are available on this GitHub repo: https://github.com/alejoe91/multimodalfitting. Finally, the raw electrophysiology data (simultaneous patch-clamp and extracellular HD-MEA recordings) are available in Neurodata Without Borders (NWB) format [65, 61] on the DANDI archive: https://dandiarchive.org/dandiset/000294.

## Acknowledgement

This study was supported by the ETH Zurich Postdoctoral Fellowship 19-2 FEL-17 (APB), the ERC Advanced Grant 694829 “neuroXscales” (JB, XX, TG, VE, AH), the China Scholarship Council (XX), by funding to the Blue Brain Project, a research center of the École polytechnique fédérale de Lausanne (EPFL), from the Swiss government’s ETH Board of the Swiss Federal Institutes of Technology (TD, DM, MZ, AJ, HM, WVG), and by the European Union’s Horizon 2020 Framework Programme for Research and Innovation under the Specific Grant Agreement No. 945539 (Human Brain Project SGA3) (TD, AJ).

---

**Supplementary Figure 1:**
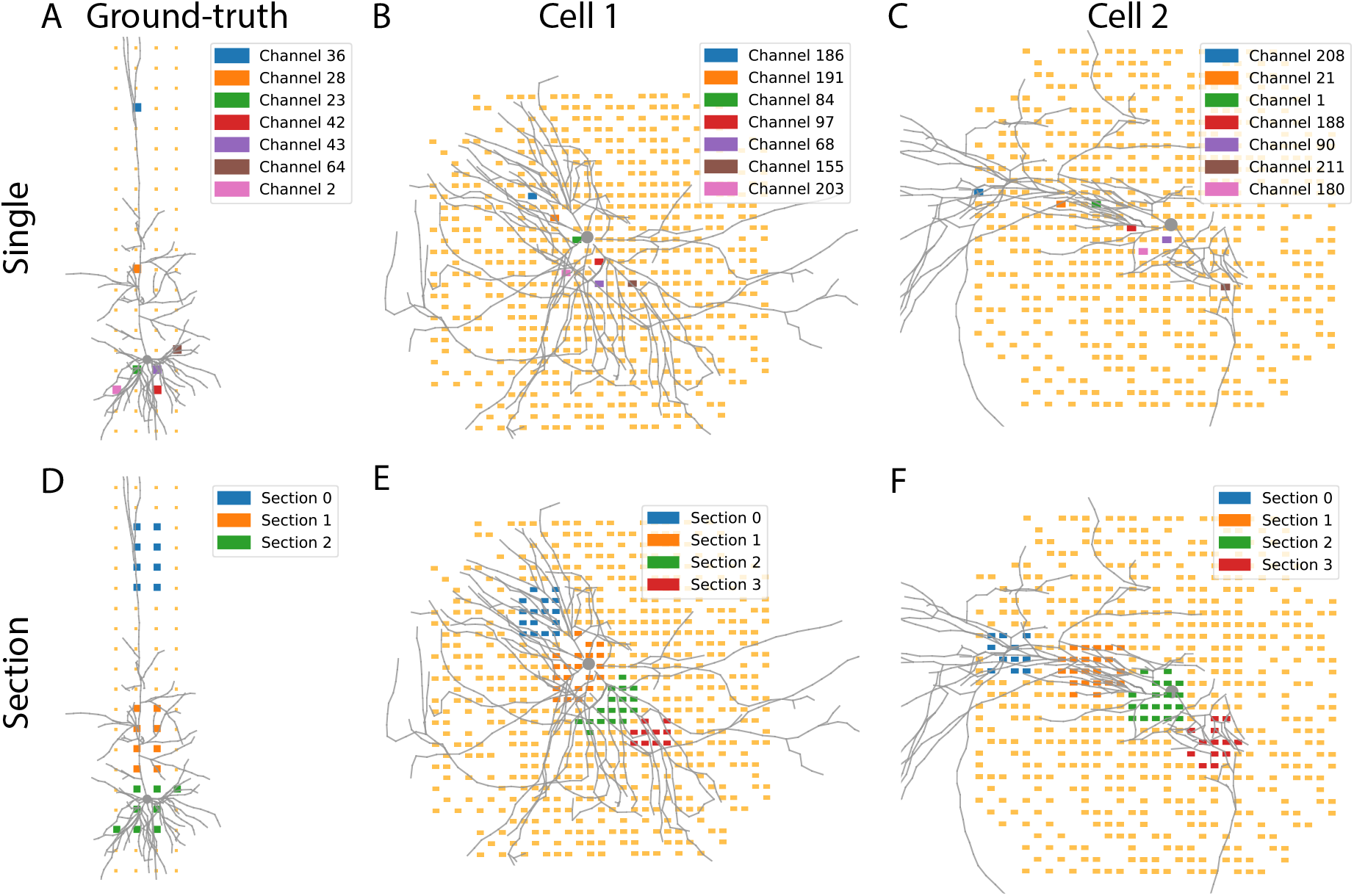
Channel selection for *single* and *sections*. Visualization of channels selected for the *single* strategy (top row) and *sections* strategy (bottom row) for the ground-truth (A-D), Cell 1 (B-E) and Cell 2 (C-F) models.

